# SpaCEy: Discovery of Functional Spatial Tissue Patterns by Association with Clinical Features Using Explainable Graph Neural Networks

**DOI:** 10.64898/2025.12.12.693857

**Authors:** Ahmet Sureyya Rifaioglu, Egle Helene Ervin, Ahmet Sarigun, Deniz Germen, Bernd Bodenmiller, Jovan Tanevski, Julio Saez-Rodriguez

## Abstract

Tissues are complex ecosystems tightly organized in space. This organization influences their function, and its alteration underpins multiple diseases. Spatial omics allows us to profile its molecular basis, but how to leverage these data to link spatial organization and molecular patterns to clinical practice remains a challenge. We present SpaCEy (**Spa**tial **C**linical **E**xplainabilit**y**), an explainable graph neural network that uncovers organizational tissue patterns predictive of clinical outcomes. SpaCEy learns directly from molecular marker expression by modelling tissues as spatial graphs of cells and their interactions, without requiring predefined cell types or anatomical regions. Its embeddings capture intercellular relationships and molecular dependencies that enable accurate prediction of variables such as overall survival and disease progression. SpaCEy integrates a specialized explainer module that reveals recurring spatial patterns of cell organisation and coordinated marker expression that are most relevant to predictions of the models. Applied to a spatially resolved proteomic lung cancer cohort, SpaCEy discovers distinct spatial arrangements of cells together with coordinated expression of protein markers associated with disease progression. Across multiple breast cancer proteomic datasets, it consistently stratifies patients according to overall survival, both across and within established clinical subtypes. SpaCEy also highlights spatial patterns of a small set of key protein markers underlying this patient stratification.

## 1 Introduction

Cells often organize into specific arrangements in space within tissues and the relationship between the spatial organisation of tissues and function, both in health and disease, is widely acknowledged [2, 12]. The amount of spatially resolved molecular data at the single-cell level has been rapidly increasing with technological improvements [7]. In particular, advancements in multiplex imaging enable precise mapping of cell populations and their diverse states within their tissue environment at the single-cell level, exploration of the organisation and interaction of cells in the tumour microenvironment, communication patterns, and novel disease mechanisms [1, 7]. Spatial omics also offers new translational opportunities by revealing similarities and differences in tissue organization and function among patients assigned to the same treatment group [2, 7, 46, 15, 42]. This tissue organisation frequently shows characteristic changes in spatial arrangements of cells which are shared across patients in diseases like cancer and fibrosis. Identifying these changes is critical because they are often structured as spatial motifs, which are localized structural patterns of cells and molecules within the tissue [25]. The high resolution offered by spatial omics technologies now makes it possible to identify these spatial motifs, providing an opportunity to link this spatial variation directly to clinical outcomes.

Analysing complex, high-dimensional omics data requires advanced machine learning methods capable of integrating diverse datasets, modelling relationships within and across tissues, and extracting meaningful biological insights to support the prediction of clinical outcomes. Subsequently, such models allow for studying the link between the changes in these relationships in response to perturbations, such as disease or drug treatment. Some of these methods do not incorporate spatial information for clinical outcome prediction. For example, Fast Survival SVM applies a max-margin framework for time-to-event prediction for survivability prediction [35], Random Survival Forests use ensemble tree models to capture nonlinear relationships between omics data and survival outcomes [21], and Gradient Boosting Survival Analysis builds an ensemble of decision-tree models in a stage-wise manner to capture nonlinear predictors of time-to-event outcomes [34]. In contrast, spatially informed approaches have been developed to model tissue organization directly, enabling the integration of spatial coordinates to solve complex tasks such as clinical outcome prediction and characterization of tissue microenvironments. For instance, Wu *et al*. developed a graph neural network (GNN) to model tumour microenvironments using spatial protein profiles from tissue specimens [47]. This method leverages multiplexed immunofluorescence imaging to capture the complex interactions within the tumour microenvironment and predict patient outcomes based on local cell type compositions. Similarly, STELLAR combines spatial and molecular information to enhance cell-type annotation and discovery in spatially resolved single-cell datasets. Using geometric deep learning, it assigns cells to known types, identifies novel cell states, and captures higher-order tissue structures [6]. SPACE-GM models tumour microenvironments as cellular graphs with the aim of predicting cancer recurrence and patient survival based on cell-type compositions of neighbourhoods [47]. A recent study by Ali *et al*. applied GNNs to spatial proteomics data, showing their models capture meaningful spatial features by analysing the embeddings of the samples [4]. These spatially aware methods highlight the value of incorporating tissue architecture into single-cell analysis, enabling graph neural networks to uncover deeper insights into cellular function and tissue organization.

Despite recent developments of spatially informed methods for analysing tissue organisation, major limitations remain in translating predictive power into biological or clinical insight. One of the key limitations, when applied to spatial tissue graphs, existing methods often rely on predefined cell-type labels or local clustering, which prevents the discovery of emergent organizational patterns directly from continuous molecular measurements. At the same time, the aforementioned machine learning models, despite their strong predictive performance, fall short in identifying contiguous, spatially coherent regions of importance and in effectively incorporating spatial information into their modelling frameworks [29]. They typically limit interpretability to analyses of model-derived latent representations or cell-level importance scores, thereby limiting their ability to uncover spatial motifs that are critical to clinical outcomes. Consequently, current methods lack the capacity to simultaneously revealing the spatially important regions and providing a concise set of molecular markers that collectively drive the observed clinical outcomes. Addressing these limitations is essential for improving confidence in model decisions, reliability, and the potential of obtaining novel biological insights [11], and to ultimately deploy machine learning models in clinical practice.

We present SpaCEy, an explainable method for discovering spatially contiguous important motifs associated with clinical outcomes using GNNs. Unlike existing approaches that rely on predefined cell-type labels or local clustering, SpaCEy identifies shared contiguous spatial patterns directly from measured cell-specific marker abundances. By representing each tissue sample as a spatial graph and learning predictive embeddings, SpaCEy bridges the gap between tissue architecture and clinical stratification. To overcome limitations of current approaches, which often produce predictive embeddings without revealing the structural or molecular features driving their predictions, SpaCEy integrates an explainer model that highlights spatially contiguous nodes and edges responsible for its outputs by identifying compact sub-graphs [50]. This enables interpretable insights into tissue organization and molecular determinants of disease. Furthermore, SpaCEy identifies a concise set of marker proteins that are most influential for predicting the corresponding clinical outcomes, providing molecular-level interpretability alongside spatial insights. By capturing emergent patterns across patients without requiring prior annotation, SpaCEy links complex spatial structures to clinical outcomes. By offering a scalable, explainable and data-driven framework for analysing tissue organisation, SpaCEy paves the way for novel insights into the spatial determinants of disease and their impact on patient outcomes.

## 2 Results

### 2.1 Model Architecture and Analysis Pipeline

SpaCEy is an explainable method that detects spatial patterns in tissue samples which are linked to clinical outcomes. The input to SpaCEy are samples collected from one or more independent cohorts, represented by measurements of spatially resolved omics and paired clinical observations (Fig. 1**a**). The spatially resolved samples are transformed into a spatial graph by Delaunay triangulation method [10]. This approach ensures that each cell is connected to its spatially nearest neighbours, preserving the tissue architecture in the graph structure. In these spatial graphs, each node represents a cell, and the features of each node represent the abundance of each measured protein marker in the corresponding spatial location. A GNN model is then trained on this spatial graph representation, leveraging the connectivity and spatial relationships between cells to learn meaningful latent representations of the input samples (Fig. 1**a**). To interpret the learned model, an explainer model generates feature and edge masks that highlight the most relevant components contributing to the GNN’s predictions for the clinical outcome of interest (Fig. 1**b**) [50]. The input to the explainer model consists of the trained model and constructed spatial graph, and its output is the learned edge masks (i.e. edge importance values) for the input graph. A key novelty of SpaCEy lies in its ability to generate interpretable spatially contiguous regions that uncover spatially localised molecular drivers of clinical outcomes. Node importance values are computed by aggregating edge-mask values from k-hop neighbours of each node, enabling the identification of key spatially contiguous, compact subgraphs that mark important regions in the tissue. Using these importance values together with the latent representations from the trained model, we perform unsupervised analysis of the latent space representations of samples, and perform differential analysis to extract biologically relevant insights (Fig. 1**c**). Based on the identified molecular and spatial features, SpaCEy provides a data-driven framework for understanding the heterogeneity of structure and function in the tumour microenvironment and enables an improved patient stratification (Fig. 1**c**).

**Figure 1.**
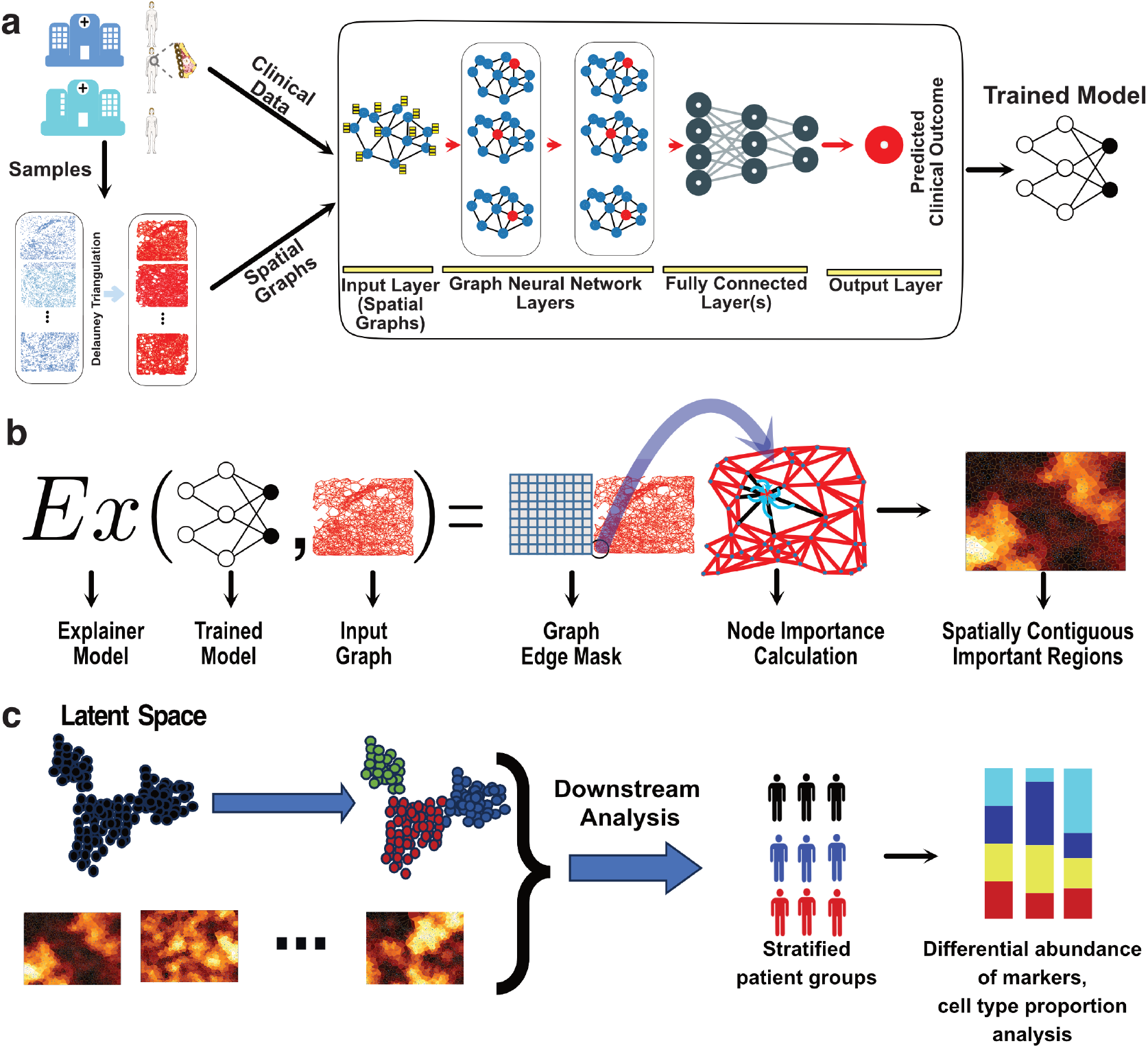
Model architecture and data analysis pipeline of SpaCEy **a)** Samples from multiple cohorts were collected as the source of high-dimensional spatial data for this study. These samples were converted into spatial graphs using the Delaunay triangulation method, and a graph neural network (GNN) was trained on these graphs, integrating spatial information and clinical data to predict relevant clinical outcomes. **b)** The trained GNN model is then analysed using the explainer model, which outputs learned edge masks. Node importance values are subsequently calculated by aggregating the edge-mask values from the k-hop neighbours of each node. These node importance values are used to identify key spatial regions within the tissue samples. **c)** Latent representations of samples and identified important regions were then used for visualization, differential analysis, and biological interpretation based on node importance values, ultimately enabling patient stratification and downstream analysis.

### 2.2 SpaCEy identifies spatially organized protein and cellular patterns predictive of tumour progression

We first applied SpaCEy to a recently published lung adenocarcinoma (LUAD) imaging mass cytometry data by Sorin *et al*. [41] comprising samples from 416 patients, with the goal of predicting tumour progression as a supervised classification task. Summary statistics and cohort characteristics of LUAD dataset is provided in Supplementary Table 1. Fig. 2a shows the distribution of progression labels across samples using a boxplot, revealing no significant difference in overall survival between the two progression groups. To investigate the relationship between the clinical outcomes and the latent representations generated by our model, we visualized the embeddings across all nodes for each sample in UMAP space (Fig. 2b), with each point representing a sample and coloured by progression status. Together, these visualizations demonstrate that the learned embedding is associated with clinical progression.

**Figure 2.**
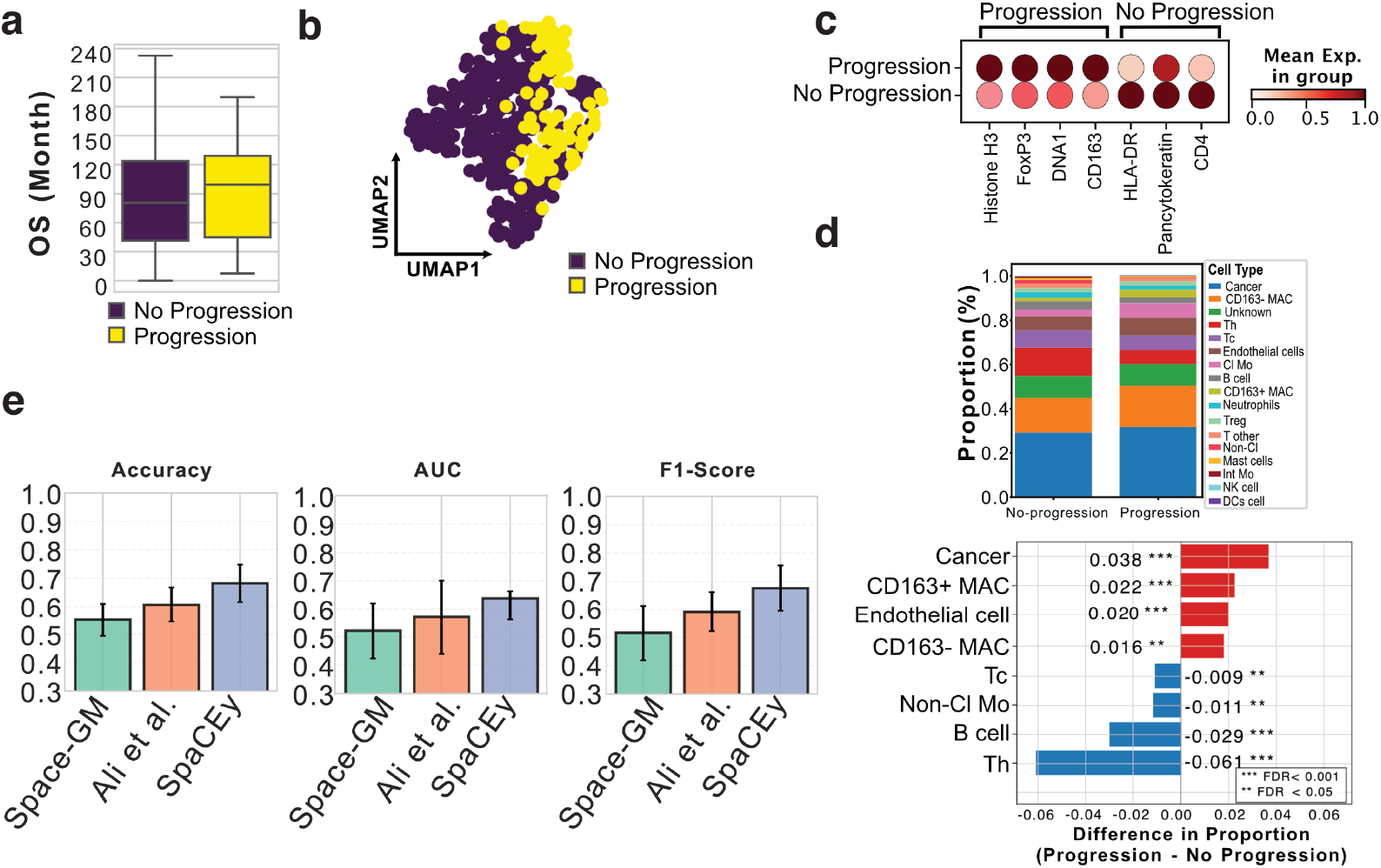
Cellular composition and progression-associated shifts in the lung tumour microenvironment. **a)** Overall survival (survival) of patients stratified by progression status, with progression and no progression **b)** A two-dimensional UMAP projection of sample-level embeddings generated from spatial proteomic features, showing the organization of samples in a low-dimensional manifold based on the clinical outcome of patients **c)** Differential abundance analysis identifying distinct protein markers between progressor and non-progressor patients. **d)** Cell type proportions of important nodes in the LUAD dataset (top panel) and significant proportional differences between progression and no-progression groups (bottom panel). Positive values indicate enrichment in the progression group, while negative values indicate enrichment in the no-progression group. **e)** Comparison of SpaCEy with the state-of-the-art methods

Analysis of marker abundance revealed distinct immune microenvironments and tumour-associated profiles between patients with and without progression Fig. 2c. Specifically, FOXP3, CD163, Histone H3, and DNA1 were enriched in the progression group. FOXP3^+^ regulatory T cells and CD163^+^ macrophages form an immunosuppressive axis previously associated with aggressive LUAD architecture and poor outcomes [41]. In contrast, non-progressing patients exhibited higher expression of HLA-DR and CD4, and a slight upregulation of epithelial cancer cell marker pan-cytokeratin. HLA-DR marks primarily antigen-presenting cells that are part of an active anti-tumour immune response [36, 30]. CD4^+^ T cells have been shown to have a higher tendency to interact with cancer cells in low-grade tumours compared to high-grade solid LUAD [41]. Therefore, SpaCEy specifically highlights these regions of cancer-immune cell interactions that are characteristic of tumours with better prognosis. Analysis of cell-type proportions revealed differences between progression and non-progression groups Fig. **2d**. Tumours from patients who progressed showed significantly increased frequencies of macrophage populations, including CD163^+^ macrophages, as well as higher cancer and endothelial cell content. This is consistent with the original publication demonstrating that CD163^+^ tumour-associated macrophages, typically linked to immunosupression and tumour progression, are enriched in high-grade tumours and strongly co-occur with FOXP3^+^ regulatory T cells [41]. The increased co-localisation and interaction between cancer and endothelial cells observed in higher-grade histological patterns reflects their aggressive nature and high metastatic potential [41]. Interestingly, SpaCEy also identifies a previously not described enrichment of CD163^−^ macrophages in regions associated with progression. In contrast, non-progressing tumours exhibited elevated numbers of Th cells, T cells and B cells. Both Th cells and CD4^+^/CD8^+^ T cells play key roles in anti-tumour immunity, whereas B cells have previously been associated with improved overall survival in LUAD [41]. Overall, these patterns indicate that SpaCEy identifies clinically relevant regions predictive of tumour progression and also shows potential to reveal novel spatial trends.

We evaluated SpaCEy against two state-of-the-art baselines, Ali *et al*. [4] and Space-GM [47], using five-fold cross-validation across all performance metrics (accuracy, F1-score, and AUC) Fig. 2e. We performed a random split stratified by disease status and fixed the splits to ensure a fair comparison and an equal distribution of disease status across folds. SpaCEy achieved the highest overall performance and the most stable behaviour across folds. In terms of accuracy, SpaCEy reached a mean of 0.68, outperforming both Ali *et al*. (0.61) and Space-GM (0.55). Similar trends were observed for F1-score, where SpaCEy got 0.68 on average, whereas the method from Ali *et al*. and Space-GM achieved 0.59 and 0.52, respectively. SpaCEy also demonstrated higher discriminative capability, yielding the highest mean AUC (0.62), compared with 0.57 for Ali *et al*. and 0.52 for Space-GM. Notably, SpaCEy consistently performed strongly across folds, including peak values of 0.77 accuracy, 0.75 F1-score, and 0.78 AUC.

### 2.3 SpaCEy stratifies breast cancer patients by subtype-agnostic localised spatial patterns

The second dataset used to train SpaCEy was an imaging mass cytometry dataset comprising 720 samples from two patient cohorts at the University Hospital of Basel and the University Hospital of Zurich [22]. The objective was to predict overall survival (hereafter referred to as survival) values using this dataset, which includes breast cancer samples across a range of clinical subtypes and cancer stages. We refer to this dataset as the JacksonFischer dataset throughout the remainder of the text. Summary statistics and cohort characteristics are provided in Supplementary Table 2. The survival values of patients grouped into different clinical subtypes show different distributions (Fig. 3**a**), where patients with triple-negative breast cancer (TNBC) exhibit the lowest survival rates due to the aggressive nature of the subtype, lack of targeted therapies, and higher likelihood of early metastasis [18, 5]. The corresponding UMAP of the embeddings learned by the SpaCEy is shown in (Fig. 3**b**). The embeddings revealed a structured distribution of samples, with similarity grouping associated primarily with survival.

**Figure 3.**
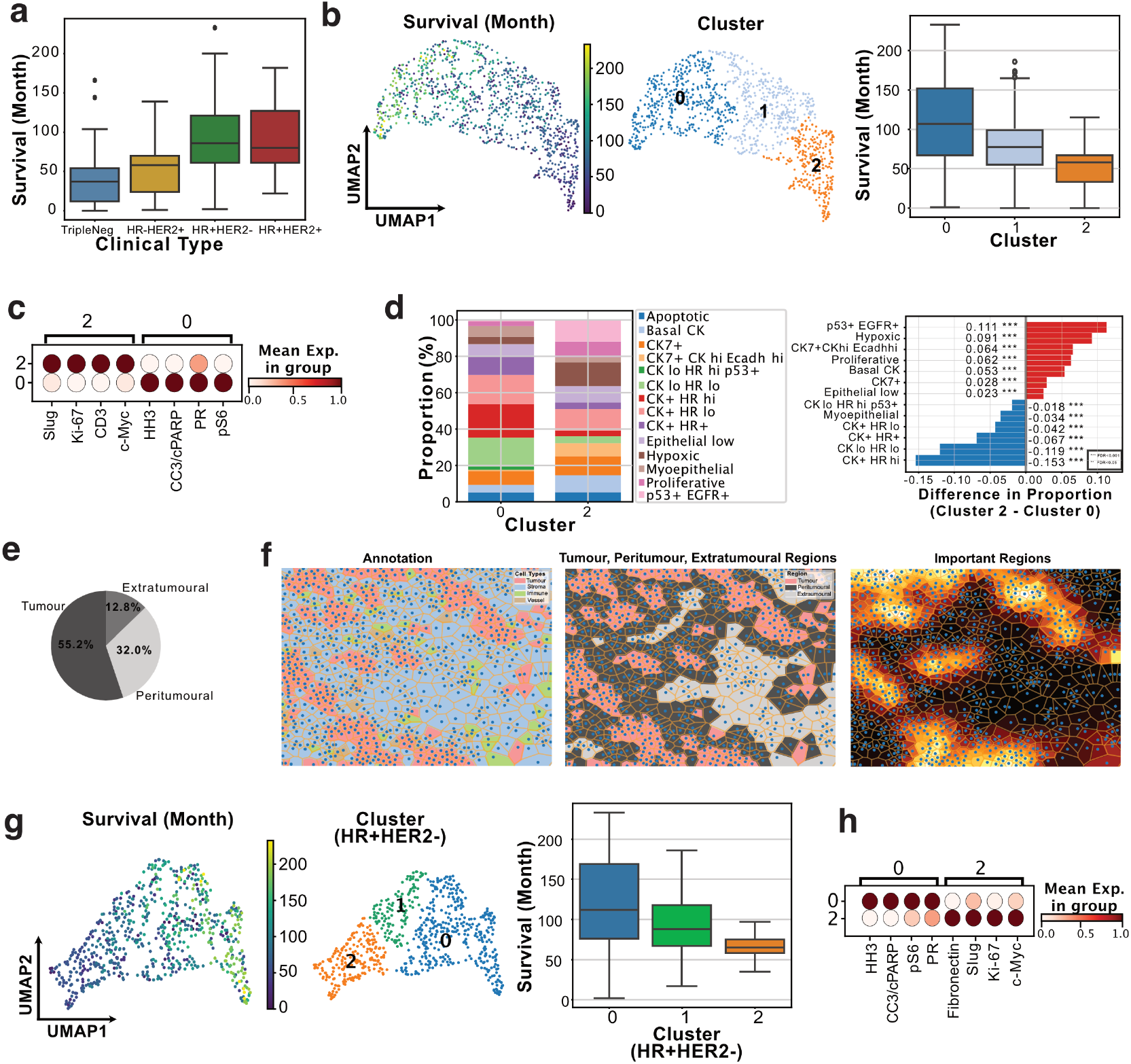
Training and analysis on JacksonFischer dataset **a)** Distribution of overall survivability values across clinical subtypes in the JacksonFischer dataset. **b)** UMAP visualization of sample embeddings derived from the trained model, showing overall survivability distributions across clusters and highlighting potential cluster-specific survivability trends. **c)** Differential abundance analysis identifying distinct protein markers between clusters, with Cluster 0 representing high-survival patients and Cluster 2 representing low-survival patients. **d)** Cell type proportions of important nodes in the JacksonFischer dataset and proportional differences between Cluster 2 and Cluster 0. Positive values indicate enrichment in Cluster 2, while negative values indicate enrichment in Cluster 0, with enrichment scores shown for each cell type. **e)** Distribution of important regions identified by the model. **f)** Cell type annotations one representative sample: first column shows overall annotation, second highlights tumour core, peritumoural, and extratumoural regions, and third indicates important regions. **g)** UMAP visualization of latent representations for HR+HER2-patients, showing clustering structure and overall survivability distributions per cluster. **h)** Differential expression analysis identifying distinct protein markers between clusters, with Cluster 0 corresponding to high-survival and Cluster 2 to low-survival patients.

We then performed unsupervised clustering based on the latent representations of the samples using the leiden algorithm [43]. Based on the resulting clusters, patients were stratified into three subtype-agnostic groups exhibiting distinct survival distributions (Fig. 3**b**). Cluster 0 was enriched for high-survival patients with high variance, showing the highest proportion of HR+HER2+ samples (45.7%) and the lowest proportion of TNBC samples (22.7%) (Fig. 3**b**, Supplementary Fig. S1). In contrast, Cluster 2 was enriched for low-survival patients, showing the highest proportion of TNBC samples (41%) and the lowest proportion of HR+HER2+ samples (12.4%). Next, we isolated the important areas identified by the explainer model, and we examined the differentially abundant proteins between these groups to investigate the differences between Cluster 2 (low-survival)- and Cluster 0 (high-survival) patients. We identified spatially-contiguous expression of protein markers distinguishing high- and low-survival clusters in the overall dataset (Fig. 3**c**). In particular, proteins associated with high-survival patients (Cluster 0) differed from those in low-survival clusters (Clusters 2). The identification of localised expression of Slug, Ki-67, CD3, and c-Myc in lower survival patients from the overall dataset suggests their association with breast cancer progression and poor prognosis. For example, Slug is a transcription factor involved in epithelial-to-mesenchymal transition (EMT), which enhances tumour invasiveness and metastasis [8]. Ki-67, a widely used proliferation marker, indicates aggressive tumour growth, and elevated Ki-67 levels are associated with poor survival in breast cancer [40, 32]. Finally, c-Myc, an oncogene involved in cell cycle regulation and proliferation, is frequently overexpressed in aggressive breast tumours and is associated with poor prognosis [48]. The combination of these markers indicates a tumour biology characterised by high proliferation (Ki-67, c-Myc), EMT-driven invasion (Slug). Notably, detecting CD3 in lower survival patients in the overall dataset suggests heterogeneity in immune cell infiltration, which could influence therapeutic responses.

To further dissect cellular composition within these key clusters, we performed cell type proportion analysis and identified the highest proportional differences based on the nodes identified as important in the JacksonFischer dataset (Fig. 3**d**. Specifically, Cluster 0 exhibited a higher abundance of hormone receptor-positive cells known to respond to endocrine therapy and, which may contribute to favourable survival [13]. In contrast, Cluster 2 was enriched with p53+ EGFR-positive, hypoxic, basal, and highly proliferative cancer cell phenotypes. Breast tumours with alteration in TP53 have a worse prognosis [14], and EGFR overactivation drives proliferation and survival signalling [31], and has been associated with poor clinical outcomes [37]. Both hypoxia and high proliferation indicate rapid tumour growth and aggressive clinical behavior [38, 51]. Basal cells, also known as myoepithelial cells, express markers of EMT and contribute to invasive behaviour [20]. Collectively, a higher proportion of cells exhibiting upregulation of oncogenic proteins, signatures and processes linked to rapid tumour growth, metastasis and therapy resistance is expected in the poor survival group. Similar associations between phenotypic composition and patient outcomes have also been reported in previous studies [17, 22]. We further characterised the localization of important regions within the tissue (Fig. 3**e**). We found that 55.2% of these regions were in tumour areas suggesting that the abundance values in the tumour areas are the most informative for the survival prediction. The second most important regions are the peritumoural areas suggesting that tumour microenvironment plays a critical role in disease progression and patient outcomes. For example, hypoxia in the tumour microenvironment can lead to genetic instability and more aggressive tumour phenotypes. In breast cancer specifically, hypoxic regions are linked to metastasis, treatment resistance, and poor prognosis and these factors may contribute to the observed spatial distribution of important regions and their association with patient survival [27, 28]. An example case is given in Fig. 3**f** where these important regions predominantly align with the tumour core and peritumoural areas.

### 2.4 SpaCEy explains the heterogeneity of the survival of patients within a clinical subtype

Hormone receptor-positive, HER2-negative (HR+HER2-) breast cancer represents the most prevalent molecular subtype, accounting for the majority of breast cancer diagnoses. Although patients with HR+HER2-tumours generally exhibit more favourable prognoses compared to more aggressive subtypes such as TNBC, substantial heterogeneity remains in disease progression, therapeutic response, and overall clinical outcomes [16]. This variability presents a significant challenge for precision oncology, as current classification systems may not adequately capture the biologically relevant distinctions within this subgroup. To address this gap, we investigated the latent molecular architecture of HR+HER2-tumours using spatially resolved proteomic data. As shown in the UMAP embedding, patient samples with similar survival tend to cluster together, and the survival distributions across clusters reveal marked differences, specifically, reduced survival in cluster 2 and higher survival values in cluster 0 in Fig. 3**g**. Notably, samples within HR+HER2-subtype show varying survival, underscoring the biological heterogeneity that exists within this molecular subtype and can be group into three clusters.

Beyond patient re-stratification, SpaCEy allows for improved insights into previously well-established clinical subtypes and heterogeneity of outcome. When comparing explainer features between the entire dataset and the HR+HER2-subgroup (Fig. 3**h**), we observed a significant overlap in the survival-associated proteins. The high expression of Ki-67, Slug, and c-Myc in the lower-survival samples of both the whole data set and HR+HER2-subset highlights the central role of these proteins in tumour proliferation and progression in breast cancer in general. The elevated expression of fibronectin in the lower-survival HR+HER2-patients points to variations in metastatic potential and epithelial characteristics [39]. More broadly, we observed that high expression of apoptotic markers, such as cleaved caspase and PARP, was enriched in the high survival group. The spatial expression patterns of these proteins enhance our comprehension of the biological diversity within HR+HER2-breast cancer, and can be potentially used as a guide to personalised treatment strategies.

### 2.5 SpaCEy generates generalizable insights across cohorts

We used the METABRIC dataset as a second independent breast cancer cohort to train a separate model [3]. This allowed us to further investigate the relationship between patient survival and spatially resolved proteomic features. The METABRIC dataset comprises 460 imaging mass cytometry samples from 405 patients, capturing diverse clinical features. The preprocessing steps are described in the Data and Preprocessing subsection, and summary statistics of the METABRIC dataset are presented in Supplementary Table 3. We calculated the survival distributions across different clinical subtypes (Fig. 4**a**). Similar to our previous observations, UMAP embeddings of the METABRIC dataset (Fig. 4**b**) demonstrated relevant patterns associated with overall survivability, with Cluster 0 being predominantly associated with high survival and Cluster 2 with lower survival. Differential protein abundance analysis identified distinct molecular markers distinguishing these survival-associated clusters (Fig. 4**c**). In Cluster 0 (higher survival), we observed enrichment of HER2 and CK8/18. Conversely, Cluster 2 (lower survival) was characterised by increased expression of fibronectin, vimentin, beta-catenin, and SMA. Consistent with prior findings in JacksonFischer dataset, elevated fibronectin expression has been linked to poor prognosis and is associated with increased invasiveness and metastatic potential [26]. A high abundance of vimentin, a well-known marker of EMT, is associated with enhanced migratory and invasive behaviour in breast cancer cells, promoting metastasis. This observation is consistent with the clinical characteristics of patients within this cluster [49]. Notably, PR and PanCK were highly expressed across the two clusters, with PR showing relatively higher abundance in the higher survival group, suggesting a shared molecular signature among patient subgroups. The occurrence of these patterns across datasets indicates that the SpaCEy captures underlying biological signals that are conserved across cohorts, despite differences in molecular panels.

**Figure 4.**
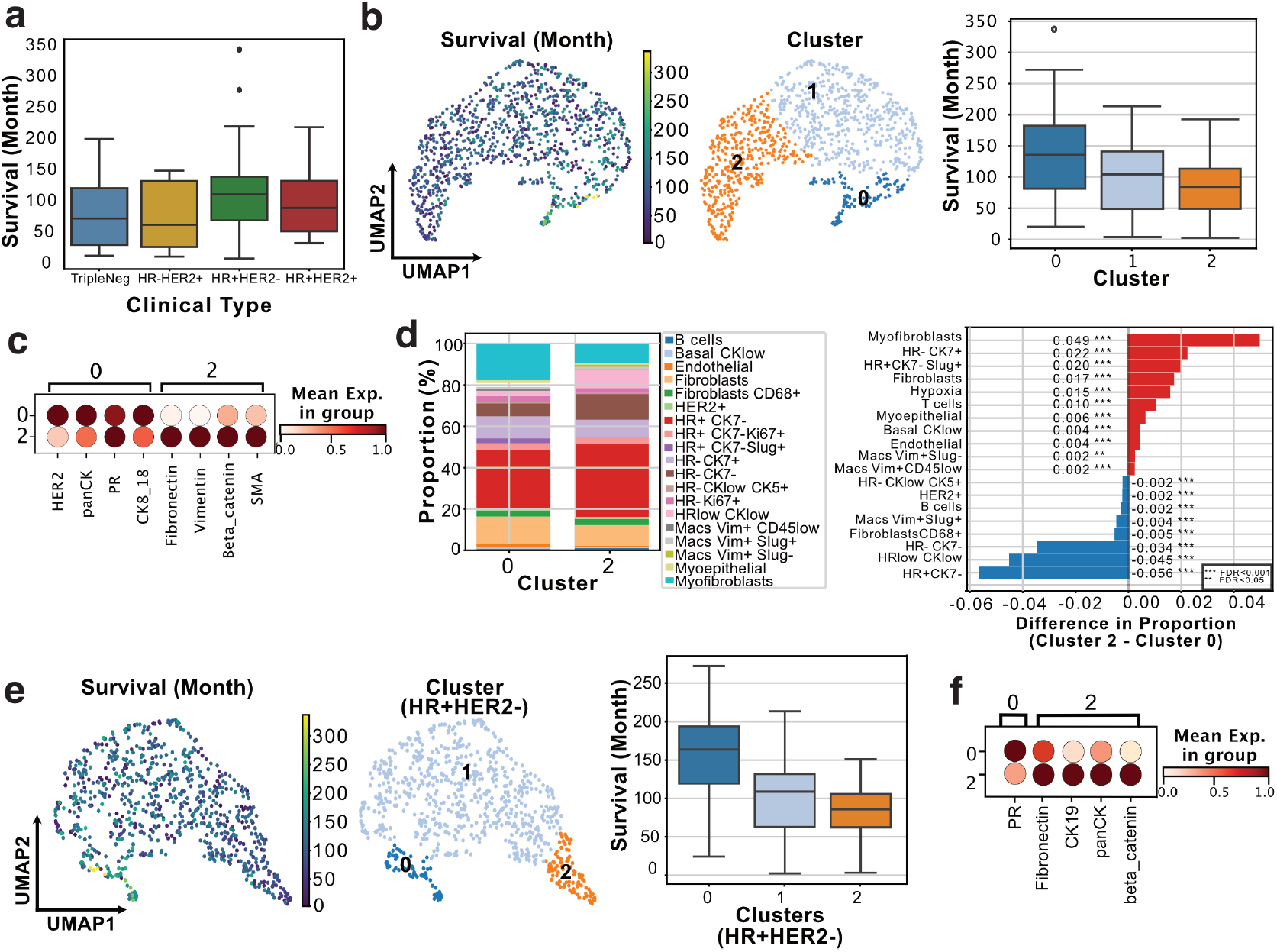
**a)** Distributions of overall survivability values of clinical subtypes **b)** UMAP visualization of patient samples colored by survival values and clustering results along with the distribution of survival values across different clusters **c)** Marker expression profiles across lower- and higher survival clusters **d)** Cell type proportions of important nodes in the METABRIC dataset and cell type proportion differences between Cluster 2 and Cluster 0. **e)** UMAP visualization of HR+HER2-samples colored by survival values and clusters and distribution of survival values across different clusters in HR+HER-samples **f)** Differential expression analysis identifies distinct protein markers between clusters, with Cluster 0 representing high-survival patients and Cluster 2 representing low-survival patients in the dataset.

To further dissect the cellular composition underlying these clusters, we compared the proportions of cell types across clusters and calculated significant cell type proportional differences between clusters (Fig. 4**d**). As in JacksonFischer dataset, we observed a higher proportion of hormone receptor-positive cells in Cluster 0, while hypoxic and myoepithelial/basal were more abundant in Cluster 2, which is associated with worse survival outcomes. The METABRIC dataset also allowed analysis of tumour microenvironment and revealed a higher proportion of myofibroblasts and fibroblasts in patients with lower survival, demonstrating the role of fibrotic stroma and cancer-associated fibroblasts in promoting malignant growth and tumour invasion [19]. Focusing on the HR+HER2-clinical subtype, the visualisation of the learned latent representations in UMAP space (Fig. 4**e**) reveals clustering of samples according to patients’ overall survival profiles. The distribution of overall survival across these clusters is consistent with our previous observations, with Cluster 2 enriched for patients with shorter survival and Cluster 0 comprising those with longer survival times. Next, we explored differential protein abundance across HR+HER2-patient clusters (Fig. 4**f** ). In Cluster 0 (higher survival), we observed higher localized expression of PR while Cluster 2 (lower survival) showed an enrichment of fibronectin, *β*-catenin and CK19. Higher abundance of fibronectin in the lower survival group is consistent with the overall analysis above. Increased *β*-catenin expression in breast cancer is significantly associated with poor prognosis, as demonstrated by a study where patients exhibiting nuclear and/or cytoplasmic *β*-catenin expression had reduced disease-specific survival rates [45]. The association between PR (progesterone receptor) expression and favourable prognosis in hormone receptor-positive breast cancer is widely acknowledged. A meta-analysis demonstrated that patients with high PR expression had significantly better survival compared to those with low PR expression [52]. These results highlight partially overlapping spatial and molecular patterns in the METABRIC and Jackson-Fischer datasets, reinforcing the association between protein abundance, tumour cellular composition and patient survival outcomes, and demonstrating SpaCEy’s ability to consistently identify prognostic patterns.

### 2.6 SpaCEy improves prediction performance using spatially resolved data

We also benchmarked SpaCEy against other machine-learning methods that do not incorporate spatial information, using the JacksonFischer and METABRIC datasets. This comparison enabled a direct assessment of whether spatial context improves predictive performance. To this end, we compared our results with three related methods: Fast Survival SVM [35], Random Survival Forest [21], and Gradient Boosting Survival Analysis [34]. To perform this comparison, we generated pseudobulk profiles of the samples based on protein abundance values and cell type annotations. For protein abundance, we applied four aggregation functions (minimum, maximum, mean, and sum) resulting in four distinct training datasets. For cell type-based aggregation, we created two additional training datasets using mean and sum aggregators (Fig. 5**a**).

**Figure 5.**
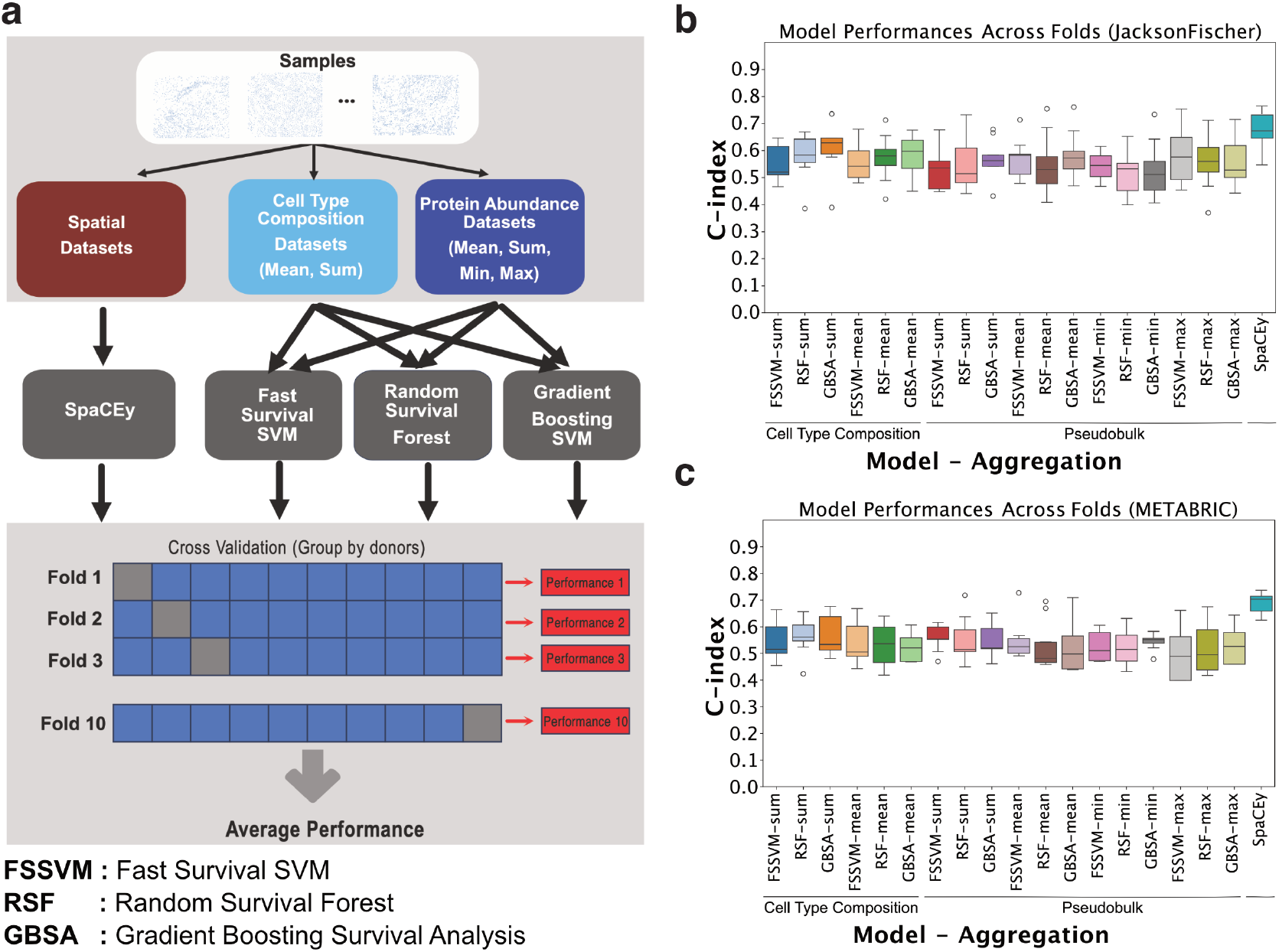
Comparison with the baseline methods a) Baseline comparison pipeline b) Performance comparison on JacksonFischer Dataset 4c) Performance comparison on METABRIC Dataset

We performed a comprehensive hyperparameter optimisation for each method-aggregation combination (Supplementary Table 4), optimising a total of 18 models. To ensure methodological rigor and a fair comparison across models, we employed consistent cross-validation splits in which folds were stratified at the patient level to prevent data leakage. The results for the JacksonFisher and METABRIC datasets are presented in Fig. 5**b** and Fig. 5**c**, respectively. Model performance was evaluated using the concordance index (C-index) as the primary metric. The choice of aggregation functions was motivated by their ability to capture different aspects of the protein abundance values, while mean aggregation smooths out individual variations, maximum and minimum values preserve extreme signals that might be biologically relevant. Sum aggregation, on the other hand, retains total protein abundance levels, which could be important for capturing global trends in tumour heterogeneity.

On the Jackson-Fischer dataset, SpaCEy achieved a mean performance of 0.720 ± 0.061, significantly outperforming all 18 baseline methods evaluated through 10-fold cross-validation. The best-performing baseline method was Cell Type Composition with Gradient Boosting SurvivalAnalysis using sum aggregation, which achieved 0.617 ± 0.092. SpaCEy demonstrated a substantial improvement of 0.102, representing a 16.5% relative improvement over this best baseline method.

On the METABRIC dataset, SpaCEy achieved a mean performance of 0.688 ± 0.038, again significantly outperforming all baseline methods. The best-performing baseline method was Cell Type Composition with Gradient Boosting Survival Analysis using sum aggregation, which achieved 0.570 ± 0.071. SpaCEy demonstrated an even larger improvement of 0.118, representing a 20.7% relative improvement over this best baseline method. The lower standard deviation of SpaCEy’s performance compared to baseline methods indicates more stable and reliable predictive capability across both datasets.

Importantly, our proposed model demonstrated the highest and most robust performance, with a clear improvement in median C-index compared to all alternative methods in both datasets. This gain was particularly evident in the METABRIC cohort, where traditional models showed more variable performance. These results indicate that our model generalizes well across cohorts and outperforms existing approaches by leveraging aggregated cellular features more effectively.

### 2.7 Scalability Analysis Results

To ensure our model remains computationally practical for large-scale spatial datasets, we assessed the computational scalability of the model across 30 configurations varying in both marker dimensionality and sample size (Fig. 6). The benchmarking workflow comprised four main stages: synthetic spatial data generation, model training and simulation, computation of performance metrics, and downstream analysis (Fig. 6**a**). This systematic analysis enabled quantitative evaluation of runtime and GPU memory usage under diverse experimental conditions. The complete results of scalibility analysis are given in Supplementary Table 5.

**Figure 6.**
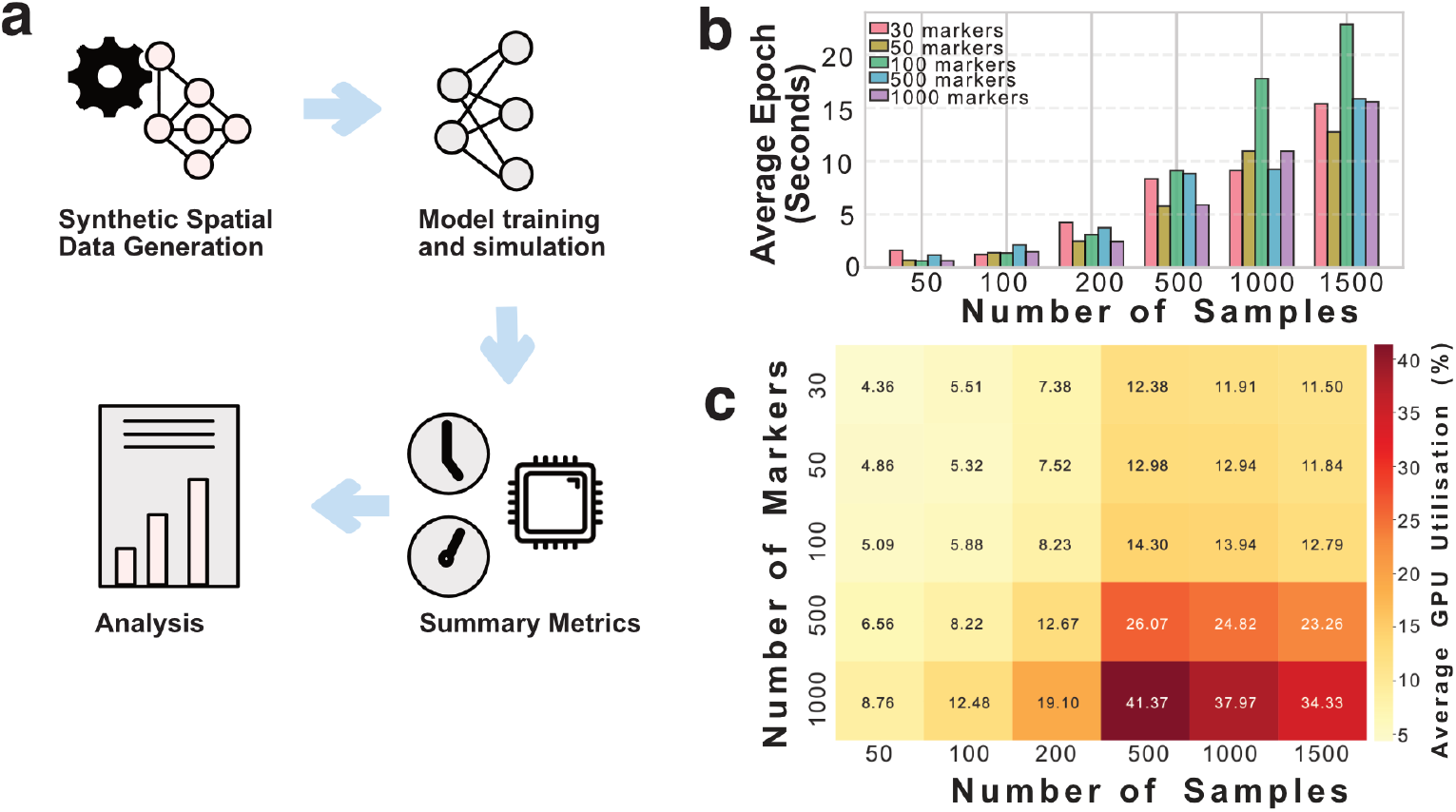
Benchmarking computational performance for synthetic spatial modelling workflows. a) Overview of the computational pipeline comprising synthetic spatial data generation, model training and simulation, computation of summary metrics, and downstream analysis. b) Average epoch time (in seconds) as a function of the number of samples and markers, showing the scaling behaviour of training time with increasing dataset size. c) Heatmap of average GPU memory utilization (%) across varying numbers of samples and markers, highlighting resource demands for large-scale simulations.

Model optimization was performed over multiple epochs, where one epoch corresponds to a complete pass of the model through the entire training dataset. Each epoch therefore represents a single iteration of parameter updates across all samples. Average epoch time ranged from 0.63 to 22.86 seconds (mean = 6.89 ± 6.12 s), highlighting favourable performance across all tested conditions (Fig. 6**b**). For small datasets (50 samples), all marker configurations exhibited comparable performance (0.95 ± 0.43 s per epoch), enabling rapid model prototyping. Epoch time increased approximately linearly with dataset size, reaching 7.59 ± 1.63 s at 500 samples, 11.60 ± 3.56 s at 1000 samples, and 16.50 ± 3.77 s at 1500 samples. Interestingly, a non-monotonic trend was observed between marker count and training time, with intermediate dimensionality (100 markers) resulting in slower performance at large sample sizes (20.32 s vs. 12.48 s for other configurations with ≥ 1000 samples), potentially due to suboptimal GPU memory access patterns. Overall, the model exhibited sub-linear scaling behaviour, with persample processing time decreasing at larger batch sizes, suggesting improved computational efficiency for large-scale, population-level analyses.

GPU memory utilization remained modest across all configurations, ranging from 4.36 % to 41.37 % (mean = 14.14 ± 9.90 %), with a maximum absolute memory usage of only 235.18 MB for the largest configuration (1000 markers, 500 samples) (Fig. 6**c**). Memory consumption increased primarily with marker dimensionality (30 markers: 8.84 %; 1000 markers: 25.67 %) rather than sample size, reflecting the feature-centric architecture of the model. Notably, memory efficiency improved by 9.7-fold at larger batch sizes (0.072 MB/sample at 1500 samples vs. 0.701 MB/sample at 50 samples), confirming that the model scales favourably in both compute time and memory usage.

## 3 Discussion

SpaCEy introduces an explainable graph neural network method for spatial omics analysis that identifies tissue regions that are linked to clinical outcomes. Although graph neural networks have shown considerable success in biomedical prediction tasks, most existing methods largely lack mechanisms to capture spatially organised tissue patterns or to isolate a concise set of informative markers. SpaCEy addresses this gap through an integrated explainer module that highlights the contiguous spatial regions and molecular features most responsible for predictions. The resulting explanations provide interpretable representations of the tumour microenvironment, revealing patterns that distinguish patients with different outcomes. This capability moves beyond pure prediction, offering a way to generate hypotheses about how tissue structure and cellular interactions influence disease progression and patient survivability. In summary, SpaCEy enables patient stratification through analysis of learned representations while simultaneously translating these insights into spatially resolved marker profiles by capturing organizational spatial patterns of a focused set of protein markers.

We evaluated SpaCEy across three datasets and two different predictive settings. First, we applied it to a lung adenocarcinoma (LUAD) imaging mass cytometry dataset in a classification setting to predict tumour progression. Second, we trained our method on two breast cancer datasets, where SpaCEy identified survival-associated motifs while also capturing dataset-specific differences. These results indicate that it can detect both generalizable biological signals and context-dependent variations. Although demonstrated here on lung and breast cancer cohorts, the design of SpaCEy is broadly applicable and can be extended to other disease contexts. Importantly, its ability to link structural tissue motifs to clinical outcomes highlights the potential of SpaCEy as a discovery tool across a wide range of diseases.

We compared our method with two state-of-the-art spatially aware methods, and we showed that it achieved better performance on different metrics while identifying spatially important regions and markers [4, 47]. We also compared SpaCEy with well-established non-spatial methods to assess its relative performance and to demonstrate that explicitly incorporating spatial information is essential for performance improvement and for capturing biologically meaningful spatial structure [21, 35, 34]. We demonstrated a clear performance improvement over the non-spatial methods, which often rely on aggregated features that obscure spatial context, limiting their ability to capture cell–cell dependencies. At the same time, unlike many graph-based methods that prioritize predictive accuracy at the expense of transparency, SpaCEy balances performance with explainability. This combination is particularly valuable in biomedical research, where the interpretability of machine learning models is critical for facilitating translation into clinical settings.

This study also has limitations. The current study focuses on protein abundance data, which provides only a partial view of cellular states and interactions. As broader multimodal datasets such as transcriptomic or metabolomic measurements become more widely accessible, and spatial technologies scale to larger tissue sections and higher molecular resolution, computational efficiency will become increasingly important. Future developments of SpaCEy will therefore incorporate strategies for scalable graph training to better accommodate the rapidly growing size and increasing complexity of multi-modal spatial datasets. Additionally, the quality of explanations is tied to the performance of the predictive models, therefore, the explanations based on the modest performing models should be interpreted conservatively. Moreover, similar to the most feature-importance–based interpretability methods, SpaCEy identifies features and spatial motifs that are important for prediction, not necessarily causal determinants of an outcome. Recognising these limitations is critical for avoiding overinterpretation of explanatory outputs and for guiding future work toward more rigorous causal interpretation frameworks.

In conclusion, SpaCEy represents a step toward explainable machine learning for spatial omics. By combining predictive accuracy with transparent explanation maps, it provides a framework that not only models clinical outcomes but also generates biologically interpretable insights into tissue structure. Its flexibility, scalability, and focus on explainability make SpaCEy a promising approach to advance spatial biology and precision oncology. As spatial omics technologies continue to evolve, SpaCEy can serve as a foundation for bridging high-dimensional molecular data with clinically actionable understanding of disease.

## 4 Methods

### 4.1 Data and preprocessing of the data

Transforming input data into a structured format is a key first step in developing graph-based algorithms for predictive modelling. For our datasets, we performed a transformation of the input data into spatial graphs using Delauney triangulation method, where each node represents a cell and the features correspond to protein abundance values [10]. These spatial graphs were then used as input for machine learning methods to address predictive problems.

We used three datasets in this study. The first dataset comprises samples from 416 patients with lung adenocarcinoma [41]. In this dataset, a 35-plex antibody panel was optimised to identify cancer cells, stromal cells, and innate and adaptive immune lineages with diverse functional substates. The second is the JacksonFischer dataset, comprising 720 imaging mass cytometry samples from 352 breast cancer patients across two cohorts: the University Hospital of Basel and the University Hospital of Zurich. This dataset also includes clinical features such as tumour size, tumour grade, clinical subtype, age, and overall survival [22]. The third dataset is the METABRIC dataset, which consists of 460 images from 405 unique patients [3]. Both datasets include clinical features of patients such as tumour size, tumour grade, clinical subtype, age, and overall survival. The basic statistics about these datasets are available in Supplementary Table 1, 2 and 3, respectively.

The preprocessing pipeline transforms spatial proteomics data into graph neural network-ready format through a four-step process. First, data quality filtering removes images with fewer than 100 cells to ensure sufficient spatial density for meaningful analysis. Next, an adaptive edge length distribution calculation is performed for each sample. To reduce spurious long-range connections introduced by the graph construction process, edges longer than the 99th percentile of all edge lengths were removed. The graph construction phase then generates spatial graphs by applying Delaunay triangulation and retaining only edges shorter than the adaptive threshold, effectively filtering out long-distance connections that may not represent biologically relevant edges. As a data augmentation step, we implemented a patch-based approach to enhance computational efficiency and increase the number of training samples. Specifically, images with cell counts exceeding the median were divided into four spatial quadrants (upper-left, upper-right, lower-left, and lower-right) while maintaining their original spatial structure. Finally, the data preprocessing pipeline extracts and serializes node features (protein abundance values), edges, clinical metadata (grade, tumour size, treatment, progression, survival data), and cell type annotations.

### 4.2 Model Architecture

We developed a model architecture, illustrated in Fig. 1, consisting of graph convolutional layers followed by fully connected (feed-forward) layers. The output layer consists of a single neuron that predicts a continuous value. After each layer, batch normalisation is applied preceding the ReLU activation function [33]. We used three different GNN layers to create predictive models, which are Graph Convolutional Neural Networks (GCN) [24], Graph Attention Networks [44], and Principal Neighbourhood Aggregation [9]. We created a generic framework that allows flexible training using a choice of GNN architecture. The hyperparameters used for model training, along with their corresponding value ranges, are provided in the Supplementary Table 6.

To project cell features and their relational structure into a new embedded space, the node features 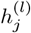 from layer *l* are first passed through a non-linear transformation function *ψ*, which in this case is the sigmoid function. The aggregation of neighbouring information is determined by the graph’s topology. This topology is represented by a normalized adjacency matrix *c*_*ij*_, which encodes the spectrally normalized edge weights between nodes *i* and *j*.

Following this transformation, an aggregation operation is applied across neighbouring nodes. Equation (1) summarizes one of the message-passing mechanism employed in our models and provides the expression of a graph convolutional network (GCN) layer.The aggregated features from neighbouring nodes within the *l*-hop neighbourhood are then concatenated with the current layer representation of node *i*, and the result is modulated by a learnable global weight parameter. This process produces the representation of node *i* at the next layer, *l* + 1. The resulting layer can be formally expressed as shown in Equation (1).

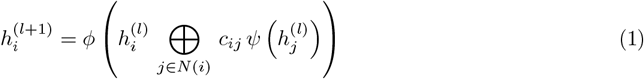

Here, 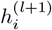 represents the updated hidden representation of node *i* at layer *l* + 1, while 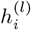 denotes its representation at the previous layer *l*. The symbol *ϕ* refers to a learnable global weight parameter, and *ψ* is a non-linear activation function. The term 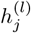 corresponds to the hidden features of neighbouring nodes *j*, and *c*_*ij*_ indicates the spectrally normalized adjacency matrix element that modulates the influence of node *j* on node *i*. Finally, ⊗ denotes the aggregation operator (e.g., max, mean, or sum) used to combine neighbourhood information.

The loss function for the survival prediction used in this study is adapted from the DeepSurv model [23], which applies a regularised negative log partial likelihood to handle survival data consisting of three components: a patient’s baseline features *x*, the event or censoring time *T*, and an event indicator *E* that specifies whether the event occurred. The objective of the model is to minimise (2):

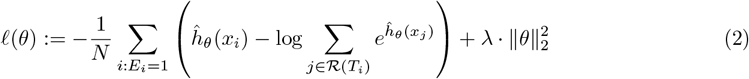

where *N* denotes the number of individuals in the cohort, *ĥ*_*θ*_ (*x*_*i*_) is the predicted log-risk for an individual *i*, ℛ (*T*_*i*_) denotes the risk set comprising all individuals still at risk at time *T*_*i*_, *E*_*i*_ ∈ {0, 1} indicates whether the event was observed, and *λ* is the coefficient of the *L*_2_ regularisation term.

For the classification tasks, we used *binary cross-entropy* as the loss function which measures the discrepancy between the predicted probability of the positive class and the true binary label. It is defined as:

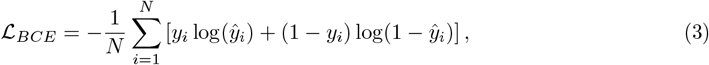

where *y*_*i*_ ∈ {0, 1} is the true label for sample *i*, and *ŷ*_*i*_ is the predicted probability of the positive class.

### 4.3 Training, Validation and Test Settings

To ensure robust model evaluation, we performed cross-validation while accounting for the presence of multiple samples from the same patients. To prevent data leakage and avoid overestimating model performance, samples originating from the same patient were grouped within the same fold. This ensured that the training and test sets remained independent with respect to patient identity. The same splits were used across all models to enable a fair comparison. We optimised hyperparameters separately for each method-aggregation combination using an internal cross-validation procedure within the training set.

### 4.4 Explainer Model

Here, we adapted the GNNExplainer method, a perturbation-based approach designed to quantify the sensitivity of model predictions to input perturbations, thereby facilitating the interpretation of graph neural network decisions. This method learns two types of masks: a discrete feature mask *M*_*F*_ over node features and a continuous edge mask *M*_*E*_, both of which identify the most influential components in the graph through an optimisation process (Supplementary Fig. S2).

The masks are initialised randomly and optimised using a loss function based on mutual information, defined in Equation (4):

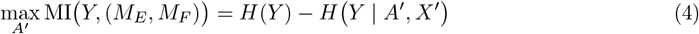

where MI denotes mutual information, *Y* represents the model predictions, *H*(·) is the entropy function, *A*^*′*^ denotes the masked adjacency matrix, and *X*^*′*^ denotes the masked node features. The mutual information is computed using the *p*_norm_ distance between two row vectors of equal dimension, as shown in Equation (5):

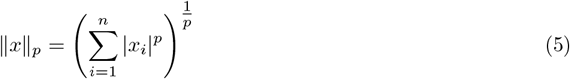

The masked features and adjacency matrix are generated as follows:

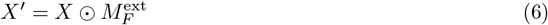

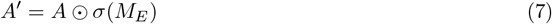

where ⊙ denotes the element-wise product and *σ* represents the sigmoid function that maps the edge mask to [0, 1]^*n×n*^.

The GNN model trained according to Equation (4) aims to maximise the mutual information between the model output *Y* and the masked subgraph. Since *H*(*Y* ) is fixed, this is equivalent to minimising the conditional entropy term. The optimisation objective is defined in Equation (8):

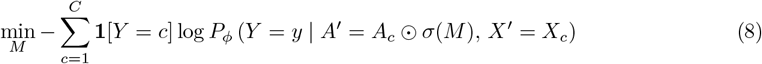

where *M* denotes the learnable mask, *X*_*c*_ the masked node features for class *c*, and *P*_*ϕ*_ the model parameterised by *ϕ*. As a result of this optimisation, the explainer model outputs the edge mask *M*_*E*_, which is subsequently used to compute node importance scores.

### 4.5 Determining the node importance values and important regions

After training the explainer model, node importance values were computed based on the learned edge weights. This process comprised the following steps:

- *Extracting k-hop subgraphs:* For each node *v*_*i*_, we defined its *k* -hop neighbourhood 𝒩_*k*_(*v*_*i*_), which includes all nodes and edges within *k* hops from *v*_*i*_.
- *Aggregating edge importance values:* Given edge importance scores *w*_*jl*_ assigned by the explainer model, we computed the node importance ℐ (*v*_*i*_) by aggregating the edge weights of all edges within the corresponding subgraph ℰ_*k*_(*v*_*i*_). Formally, the importance of node *v*_*i*_ is defined as in Equation (9):

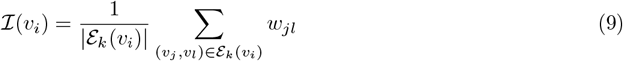

where ℰ_*k*_(*v*_*i*_) denotes the set of edges in the *k* -hop subgraph centered on node *v*_*i*_, and *w*_*jl*_ represents the importance weight assigned to the edge connecting nodes *v*_*j*_ and *v*_*l*_. The resulting node importance values were subsequently used to identify biologically meaningful regions within the tissue that contributed most strongly to the model predictions.

### 4.6 Evaluation Metrics

We assessed model performance using a comprehensive set of classification and regression metrics. For the classification task, accuracy, area under the receiver operating characteristic curve (AUC), and F1-score were used to quantify discriminative performance. Accuracy measures the proportion of correctly classified samples and is defined as:

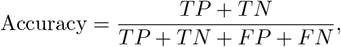

where *TP, TN, FP*, and *FN* denote true positives, true negatives, false positives, and false negatives, respectively. Area Under the ROC Curve (AUC) quantifies the model’s ability to discriminate between classes by integrating the true positive rate (TPR) as a function of the false positive rate (FPR):

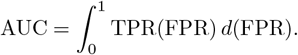

F1-score provides a harmonic mean of precision and recall and is given by:

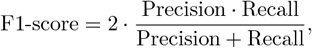

where

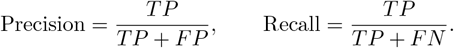

To assess the performance of our survival predictive models, we employed regression and survival evaluation metrics: Root Mean Squared Error (RMSE) and concordance index (C-index). These metrics provide complementary insights into the model’s predictive accuracy and concordance with survival outcomes. Root Mean Squared Error (RMSE) is used to measure the difference between predicted continuous outcomes and the ground truth values. It is calculated as:

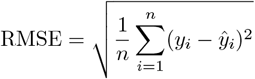

where *y*_*i*_ and *ŷ*_*i*_ represent the true and predicted values, respectively, and *n* is the number of samples. Concordance index (C-index) is a metric used in survival analysis to evaluate the model’s ability to correctly rank survival times. It is defined as:

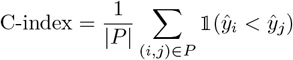

where *P* is the set of all comparable pairs, *ŷ*_*i*_ is the predicted risk score or survival time, and 𝟙_( ·) is the indicator function. A C-index of 1.0 denotes perfect concordance, while 0.5 indicates random ordering.

### 4.7 Synthetic Dataset Generation

To ensure controlled experimental conditions, we generated synthetic datasets designed to simulate the structure of real clinical tissue imaging data. Node features were represented as random vectors with dimensionality equal to the number of markers. These features were sampled from a standard normal distribution, *x* ∼ 𝒩 (0, 1), and additional noise was incorporated, *ϵ* ∼ 𝒩 (0, 0.1), to introduce variability.

For each simulation, the samples follow a normal distribution with mean 1300 and standard deviation 400, clipped between 100 and 2500. Sparse connectivity was introduced by assigning edges with probability *p* = 0.1, while ensuring overall graph connectivity. In addition, each node was assigned two-dimensional spatial coordinates to capture positional information within the simulated tissue microenvironment.

Labels were generated to provide meaningful regression targets that combined both node-level and structural information. Node importance weights were sampled from a normal distribution, *w* ∼ 𝒩 (0, 1). These were used to define a graph-level feature, *f*_graph_ = mean(*x*) · *w*, while a structural feature was derived from edge density, *f*_struct_ = |*E*|*/*(|*V* | *×* (|*V* | − 1)). The final regression label was obtained as a combination of these components with additional noise, *y* = *f*_graph_ + *f*_struct_ + *ϵ*, where *ϵ* ∼ 𝒩 (0, 0.05).

### 4.8 Scalability Analysis

To assess computational performance, we conducted a systematic scalability analysis using synthetic spatial proteomics-like datasets. The objective was to quantify how training time, GPU memory consumption, and utilization rates scale with increasing dataset dimensionality and sample size.

A factorial experimental design was implemented, varying the number of molecular markers across five levels (30, 100, 300, 500, and 1000) and the number of samples across six levels (50, 100, 200, 500, 1000, and 1500), resulting in 30 distinct configurations. For each configuration, we recorded different statistics such as average epoch training time, average GPU utilization percentage during training and memory usage. Each experiment was repeated three times to ensure consistency, and all runs were performed under identical hardware conditions. This benchmarking workflow enabled quantitative assessment of the computational scalability of our model.

## 5 Data Availability

The datasets used in this study were obtained from their original publications. The lung tumour immune microenvironment dataset was obtained from Sorin *et al*. (2023) and is publicly available from the original publication via https://doi.org/10.1038/s41586-022-05672-3. The breast cancer single-cell pathology (i.e., JacksonFischer) dataset was obtained from Jackson *et al*. (2020) and is accessible from the original publication via https://doi.org/10.1038/s41586-019-1876-x. METABRIC breast cancer dataset was obtained from Ali *et al*. (2020) and is publicly available from the original publication via https://doi.org/10.1038/s43018-020-0026-6. The preprocessed datasets used in this study are available at https://doi.org/10.5281/zenodo.17710588.

## 6 Code Availability

SpaCEy was implemented within the NVIDIA PyTorch Geometric (PyG) container (nvcr.io/nvidia/pyg:24.01-py3). The framework builds on PyTorch for deep learning and PyTorch Geometric for graph-based deep learning modelling. For downstream analysis and preprocessing, we used Scanpy (v1.9.8). In addition, Leidenalg (v0.10.2) was employed to perform Leiden clustering for community detection. All code used for data preprocessing, model training, and downstream analyses is available at https://github.com/saezlab/SpaCEy.

## 7 Hardware Environment

All computations were performed on a workstation running Ubuntu 24.04.3 LTS with Linux kernel 6.8.0-79-generic. The system was equipped with an AMD Ryzen Threadripper PRO 5995WX processor (128 cores, 2.7 GHz base frequency) and NVIDIA RTX A6000 GPU.

## 8 Author Contributions

A.S.R.: Conceptualization, Methodology, Software, Validation, Formal analysis, Data Curation, Writing – Original Draft, Writing – Review & Editing, Visualization. E.H.E.: Formal analysis, Data Curation, Writing – Review & Editing. A.S.: Methodology, Formal analysis, Software, Writing – Review & Editing. D.G.: Software, Validation, Formal analysis, Writing – Review & Editing. B.B.: Writing – Review & Editing, Supervision. J.T.: Conceptualization, Resources, Writing – Review & Editing, Supervision. J.S.R.: Writing – Review & Editing, Supervision, Funding acquisition.

## 9 Acknowledgments

ASR is supported by the Heidelberg Faculty of Medicine at Heidelberg University through the Medical Data Scientist Fellowship. AS was enrolled at the Department of Chemistry and Department of Physics at Middle East Technical University (METU) and was supported through an Erasmus Traineeship at the Institute for Computational Biomedicine, Heidelberg University and Heidelberg University Hospital, Heidelberg, Germany. DG was enrolled at the Department of Computer Engineering at Middle East Technical University (METU) and was supported through an Erasmus Traineeship at the Institute for Computational Biomedicine, Heidelberg University and Heidelberg University Hospital, Heidelberg, Germany. JT is supported by the “Bruno und Helene Jöster Stiftung”. We would like to thank Pablo Rodríguez Mier for the helpful discussions and for proofreading.

## References

[1] Method of the Year 2024: Spatial proteomics. Nature Methods, 21(12):2195–2196, December 2024. ISSN 1548-7105. doi: 10.1038/s41592-024-02565-3.

[2] Miri Adler, Arun R. Chavan, and Ruslan Medzhitov. Tissue Biology: In Search of a New Paradigm. Annual Review of Cell and Developmental Biology, 39:67–89, October 2023. ISSN 1530-8995. doi: 10.1146/annurev-cellbio-120420-113830.

[3] H. Raza Ali, Hartland W. Jackson, Vito R. T. Zanotelli, Esther Danenberg, Jana R. Fischer, Helen Bardwell, Elena Provenzano, CRUK IMAXT Grand Challenge Team, H. Raza Ali, M. A. Sa’d, S. Alon, Samuel Aparicio, G. Battistoni, S. Balasubramanian, R. Becker, Bernd Bodenmiller, E. S. Boyden, D. Bressan, A. Bruna, B. Marcel, Carlos Caldas, M. Callari, I. G. Cannell, H. Casbolt, N. Chornay, Y. Cui, A. Dariush, K. Dinh, A. Emenari, Y. Eyal-Lubling, J. Fan, E. Fisher, E. A. González-Solares, C. González-Fernández, D. Goodwin, W. Greenwood, F. Grimaldi, G. J. Hannon, O. Harris, S. Harris, C. Jauset, J. A. Joyce, E. D. Karagiannis, T. Kovačević, L. Kuett, R. Kunes, A. Küpcü Yoldaş, D. Lai, E. Laks, H. Lee, M. Lee, G. Lerda, Y. Li, A. McPherson, N. Millar, C. M. Mulvey, F. Nugent, C. H. O’Flanagan, M. Paez-Ribes, I. Pearsall, F. Qosaj, A. J. Roth, Oscar M. Rueda, T. Ruiz, K. Sawicka, L. A. Sepúlveda, S. P. Shah, A. Shea, A. Sinha, A. Smith, S. Tavaré, S. Tietscher, I. Vázquez-García, S. L. Vogl, N. A. Walton, A. T. Wassie, S. S. Watson, S. A. Wild, E. Williams, J. Windhager, C. Xia, P. Zheng, X. Zhuang, Oscar M. Rueda, Suet-Feung Chin, Samuel Aparicio, Carlos Caldas, and Bernd Bodenmiller. Imaging mass cytometry and multiplatform genomics define the phenogenomic landscape of breast cancer. Nature Cancer, 1(2):163–175, February 2020. ISSN 2662-1347. doi: 10.1038/s43018-020-0026-6.

[4] Mayar Ali, Sabrina Richter, Ali Ertürk, David S. Fischer, and Fabian J. Theis. Graph neural networks learn emergent tissue properties from spatial molecular profiles. Nature Communications, 16(1):8419, September 2025. ISSN 2041-1723. doi: 10.1038/s41467-025-63758-8.

[5] Giampaolo Bianchini, Justin M. Balko, Ingrid A. Mayer, Melinda E. Sanders, and Luca Gianni. Triple-negative breast cancer: Challenges and opportunities of a heterogeneous disease. Nature Reviews Clinical Oncology, 13(11):674–690, November 2016. ISSN 1759-4782. doi: 10.1038/nrclinonc.2016.66.

[6] Maria Brbić, Kaidi Cao, John W. Hickey, Yuqi Tan, Michael P. Snyder, Garry P. Nolan, and Jure Leskovec. Annotation of spatially resolved single-cell data with STELLAR. Nature Methods, 19 (11):1411–1418, November 2022. ISSN 1548-7105. doi: 10.1038/s41592-022-01651-8.

[7] Dario Bressan, Giorgia Battistoni, and Gregory J. Hannon. The dawn of spatial omics. Science (New York, N.Y.), 381(6657):eabq4964, August 2023. ISSN 1095-9203. doi: 10.1126/science.abq4964.

[8] Christophe Côme, Fabrice Magnino, Frédéric Bibeau, Pascal De Santa Barbara, Karl Friedrich Becker, Charles Theillet, and Pierre Savagner. Snail and slug play distinct roles during breast carcinoma progression. Clinical Cancer Research: An Official Journal of the American Association for Cancer Research, 12(18):5395–5402, September 2006. ISSN 1078-0432. doi: 10.1158/1078-0432.CCR-06-0478.

[9] Gabriele Corso, Luca Cavalleri, Dominique Beaini, Pietro Lio, and Petar Veličković. Principal Neighbourhood Aggregation for Graph Nets. In Advances in Neural Information Processing Systems, volume 33, pages 13260–13271. Curran Associates, Inc., 2020.

[10] B Delaunay. Sur la sphere vide. Bulletin of the Academy of Sciences of the USSR Classe des Sciences Mathematiques et Naturelles, 7:793–800, 1934.

[11] Finale Doshi-Velez and Been Kim. Towards A Rigorous Science of Interpretable Machine Learning, March 2017.

[12] Jun Du, Yu-Chen Yang, Zhi-Jie An, Ming-Hui Zhang, Xue-Hang Fu, Zou-Fang Huang, Ye Yuan, and Jian Hou. Advances in spatial transcriptomics and related data analysis strategies. Journal of Translational Medicine, 21(1):330, May 2023. ISSN 1479-5876. doi: 10.1186/s12967-023-04150-2.

[13] Early Breast Cancer Trialists’ Collaborative Group (EBCTCG), C. Davies, J. Godwin, R. Gray, M. Clarke, D. Cutter, S. Darby, P. McGale, H. C. Pan, C. Taylor, Y. C. Wang, M. Dowsett, J. Ingle, and R. Peto. Relevance of breast cancer hormone receptors and other factors to the efficacy of adjuvant tamoxifen: Patient-level meta-analysis of randomised trials. Lancet (London, England), 378(9793):771–784, August 2011. ISSN 1474-547X. doi: 10.1016/S0140-6736(11)60993-8.

[14] Richard M. Elledge, Suzanne A. W. Fuqua, Gary M. Clark, Pascal Pujol, and D. Craig Allred. The role and prognostic significance of p53 gene alterations in breast cancer. Breast Cancer Research and Treatment, 27(1):95–102, January 1993. ISSN 1573-7217. doi: 10.1007/BF00683196.

[15] Rong Fan. Integrative spatial protein profiling with multi-omics. Nature Methods, 21(12):2223–2225, December 2024. ISSN 1548-7105. doi: 10.1038/s41592-024-02533-x.

[16] Otto Metzger Filho, Giuseppe Viale, Shayna Stein, Lorenzo Trippa, Denise A. Yardley, Ingrid A. Mayer, Vandana G. Abramson, Carlos L. Arteaga, Laura M. Spring, Adrienne G. Waks, Eileen Wrabel, Michelle K. DeMeo, Aditya Bardia, Patrizia Dell’Orto, Leila Russo, Tari A. King, Kornelia Polyak, Franziska Michor, Eric P. Winer, and Ian E. Krop. Impact of HER2 Heterogeneity on Treatment Response of Early-Stage HER2-Positive Breast Cancer: Phase II Neoadjuvant Clinical Trial of T-DM1 Combined with Pertuzumab. Cancer Discovery, 11(10):2474–2487, October 2021. ISSN 2159-8274. doi: 10.1158/2159-8290.CD-20-1557.

[17] Jana Raja Fischer, Hartland Warren Jackson, Natalie de Souza, Zsuzsanna Varga, Peter Schraml, Holger Moch, and Bernd Bodenmiller. Multiplex imaging of breast cancer lymph node metastases identifies prognostic single-cell populations independent of clinical classifiers. Cell Reports Medicine, 4(3):100977, March 2023. ISSN 2666-3791. doi: 10.1016/j.xcrm.2023.100977.

[18] William D. Foulkes, Ian E. Smith, and Jorge S. Reis-Filho. Triple-Negative Breast Cancer. New England Journal of Medicine, 363(20):1938–1948, November 2010. ISSN 0028-4793. doi: 10.1056/NEJMra1001389.

[19] Dengdi Hu, Zhaoqing Li, Bin Zheng, Xixi Lin, Yuehong Pan, Peirong Gong, Wenying Zhuo, Yujie Hu, Cong Chen, Lini Chen, Jichun Zhou, and Linbo Wang. Cancer-associated fibroblasts in breast cancer: Challenges and opportunities. Cancer Communications (London, England), 42(5): 401–434, May 2022. ISSN 2523-3548. doi: 10.1002/cac2.12291.

[20] Mohammed Inayatullah, Arun Mahesh, Arran K Turnbull, J Michael Dixon, Rachael Natrajan, and Vijay K Tiwari. Basal–epithelial subpopulations underlie and predict chemotherapy resistance in triple-negative breast cancer. EMBO Molecular Medicine, 16(4):823–853, April 2024. ISSN 1757-4676. doi: 10.1038/s44321-024-00050-0.

[21] Hemant Ishwaran, Thomas A. Gerds, Udaya B. Kogalur, Richard D. Moore, Stephen J. Gange, and Bryan M. Lau. Random survival forests for competing risks. Biostatistics (Oxford, England), 15(4):757–773, October 2014. ISSN 1468-4357. doi: 10.1093/biostatistics/kxu010.

[22] Hartland W. Jackson, Jana R. Fischer, Vito R. T. Zanotelli, H. Raza Ali, Robert Mechera, Savas D. Soysal, Holger Moch, Simone Muenst, Zsuzsanna Varga, Walter P. Weber, and Bernd Bodenmiller. The single-cell pathology landscape of breast cancer. Nature, 578(7796):615–620, February 2020. ISSN 1476-4687. doi: 10.1038/s41586-019-1876-x.

[23] Jared L. Katzman, Uri Shaham, Alexander Cloninger, Jonathan Bates, Tingting Jiang, and Yuval Kluger. DeepSurv: Personalized treatment recommender system using a Cox proportional hazards deep neural network. BMC Medical Research Methodology, 18(1):24, February 2018. ISSN 1471-2288. doi: 10.1186/s12874-018-0482-1.

[24] Thomas N. Kipf and Max Welling. Semi-Supervised Classification with Graph Convolutional Networks, February 2017.

[25] Enikő Lázár and Joakim Lundeberg. Spatial architecture of development and disease. Nature Reviews Genetics, pages 1–19, September 2025. ISSN 1471-0064. doi: 10.1038/s41576-025-00892-5.

[26] Cheng-Lin Li, Dan Yang, Xin Cao, Fan Wang, Duan-Yang Hong, Jing Wang, Xiang-Chun Shen, and Yan Chen. Fibronectin induces epithelial-mesenchymal transition in human breast cancer MCF-7 cells via activation of calpain. Oncology Letters, 13(5):3889–3895, May 2017. ISSN 1792-1074. doi: 10.3892/ol.2017.5896.

[27] Joshua J. Li, Julia Y. Tsang, and Gary M. Tse. Tumor Microenvironment in Breast Cancer-Updates on Therapeutic Implications and Pathologic Assessment. Cancers, 13(16):4233, August 2021. ISSN 2072-6694. doi: 10.3390/cancers13164233.

[28] Yongxing Li, Fengshuo Liu, Qingjin Cai, Lijun Deng, Qin Ouyang, Xiang H.-F. Zhang, and Ji Zheng. Invasion and metastasis in cancer: Molecular insights and therapeutic targets. Signal Transduction and Targeted Therapy, 10(1):57, February 2025. ISSN 2059-3635. doi: 10.1038/s41392-025-02148-4.

[29] Zachary C. Lipton. The Mythos of Model Interpretability: In machine learning, the concept of interpretability is both important and slippery. Queue, 16(3):31–57, June 2018. ISSN 1542-7730. doi: 10.1145/3236386.3241340.

[30] Kea Martin, Jens Schreiner, and Alfred Zippelius. Modulation of APC Function and Anti-Tumor Immunity by Anti-Cancer Drugs. Frontiers in Immunology, 6, September 2015. ISSN 1664-3224. doi: 10.3389/fimmu.2015.00501.

[31] Hiroko Masuda, Dongwei Zhang, Chandra Bartholomeusz, Hiroyoshi Doihara, Gabriel N. Hortobagyi, and Naoto T. Ueno. Role of epidermal growth factor receptor in breast cancer. Breast Cancer Research and Treatment, 136(2):331–345, November 2012. ISSN 1573-7217. doi: 10.1007/s10549-012-2289-9.

[32] Sunil Sankunny Menon, Chandrasekharan Guruvayoorappan, Kunnathur Murugesan Sakthivel, and Rajan Radha Rasmi. Ki-67 protein as a tumour proliferation marker. Clinica Chimica Acta, 491:39–45, April 2019. ISSN 0009-8981. doi: 10.1016/j.cca.2019.01.011.

[33] Vinod Nair and Geoffrey E. Hinton. Rectified linear units improve restricted boltzmann machines. In Proceedings of the 27th International Conference on International Conference on Machine Learning, ICML’10, pages 807–814, Madison, WI, USA, June 2010. Omnipress. ISBN 978-1-60558-907-7.

[34] Sebastian Pölsterl. Scikit-survival: A library for time-to-event analysis built on top of scikit-learn. J. Mach. Learn. Res., 21(1):212:8747–212:8752, January 2020. ISSN 1532-4435.

[35] Sebastian Pölsterl, Nassir Navab, and Amin Katouzian. Fast Training of Support Vector Machines for Survival Analysis. In Annalisa Appice, Pedro Pereira Rodrigues, Vítor Santos Costa, João Gama, Alípio Jorge, and Carlos Soares, editors, Machine Learning and Knowledge Discovery in Databases, pages 243–259, Cham, 2015. Springer International Publishing. ISBN 978-3-319-23525-7. doi: 10.1007/978-3-319-23525-715.

[36] Paul A. Roche and Kazuyuki Furuta. The ins and outs of MHC class II-mediated antigen processing and presentation. Nature Reviews Immunology, 15(4):203–216, April 2015. ISSN 1474-1741. doi: 10.1038/nri3818.

[37] J R Sainsbury, J R Farndon, G K Needham, and A L Harris. Epidermal-growth-factor receptor status as predictor of early recurrence of and death from breast cancer. Lancet, 1(8547):1398–402, 1987.

[38] Gregg L. Semenza. The hypoxic tumor microenvironment: A driving force for breast cancer progression. Biochimica Et Biophysica Acta, 1863(3):382–391, March 2016. ISSN 0006-3002. doi: 10.1016/j.bbamcr.2015.05.036.

[39] Aparna Shinde, Sarah Libring, Aktan Alpsoy, Ammara Abdullah, James A. Schaber, Luis Solorio, and Michael K. Wendt. Autocrine Fibronectin Inhibits Breast Cancer Metastasis. Molecular Cancer Research, 16(10):1579–1589, October 2018. ISSN 1541-7786. doi: 10.1158/1541-7786.MCR-18-0151.

[40] Nahed A. Soliman and Shaimaa M. Yussif. Ki-67 as a prognostic marker according to breast cancer molecular subtype. Cancer Biology & Medicine, 13(4):496–504, December 2016. ISSN 2095-3941. doi: 10.20892/j.issn.2095-3941.2016.0066.

[41] Mark Sorin, Morteza Rezanejad, Elham Karimi, Benoit Fiset, Lysanne Desharnais, Lucas J. M. Perus, Simon Milette, Miranda W. Yu, Sarah M. Maritan, Samuel Doré, Émilie Pichette, William Enlow, Andréanne Gagné, Yuhong Wei, Michele Orain, Venkata S. K. Manem, Roni Rayes, Peter M. Siegel, Sophie Camilleri-Bröet, Pierre Olivier Fiset, Patrice Desmeules, Jonathan D. Spicer, Daniela F. Quail, Philippe Joubert, and Logan A. Walsh. Single-cell spatial landscapes of the lung tumour immune microenvironment. Nature, 614(7948):548–554, February 2023. ISSN 1476-4687. doi: 10.1038/s41586-022-05672-3.

[42] Rita Strack. Subcellular spatial proteomics. Nature Methods, 21(12):2227, December 2024. ISSN 1548-7105. doi: 10.1038/s41592-024-02546-6.

[43] V. A. Traag, L. Waltman, and N. J. van Eck. From Louvain to Leiden: Guaranteeing well-connected communities. Scientific Reports, 9(1):5233, March 2019. ISSN 2045-2322. doi: 10.1038/s41598-019-41695-z.

[44] Petar Veličković, Guillem Cucurull, Arantxa Casanova, Adriana Romero, Pietro Lio, and Yoshua Bengio. Graph Attention Networks, February 2018.

[45] Ziyi Wang, Hao Zhang, Jianxin Hou, Jianing Niu, Zhenhai Ma, Haidong Zhao, and Caigang Liu. Clinical implications of β-catenin protein expression in breast cancer. International Journal of Clinical and Experimental Pathology, 8(11):14989–14994, 2015. ISSN 1936-2625.

[46] Johannes Wirth, Nina Huber, Kelvin Yin, Sophie Brood, Simon Chang, Celia P. Martinez-Jimenez, and Matthias Meier. Spatial transcriptomics using multiplexed deterministic barcoding in tissue. Nature Communications, 14(1):1523, March 2023. ISSN 2041-1723. doi: 10.1038/s41467-023-37111-w.

[47] Zhenqin Wu, Alexandro E. Trevino, Eric Wu, Kyle Swanson, Honesty J. Kim, H. Blaize D’Angio, Ryan Preska, Gregory W. Charville, Piero D. Dalerba, Ann Marie Egloff, Ravindra Uppaluri, Umamaheswar Duvvuri, Aaron T. Mayer, and James Zou. Graph deep learning for the characterization of tumour microenvironments from spatial protein profiles in tissue specimens. Nature Biomedical Engineering, 6(12):1435–1448, December 2022. ISSN 2157-846X. doi: 10.1038/s41551-022-00951-w.

[48] Jinhua Xu, Yinghua Chen, and Olufunmilayo I. Olopade. MYC and Breast Cancer. Genes & Cancer, 1(6):629–640, June 2010. ISSN 1947-6027. doi: 10.1177/1947601910378691.

[49] Nami Yamashita, Eriko Tokunaga, Hiroyuki Kitao, Yuichi Hisamatsu, Kenji Taketani, Sayuri Akiyoshi, Satoko Okada, Shinichi Aishima, Masaru Morita, and Yoshihiko Maehara. Vimentin as a poor prognostic factor for triple-negative breast cancer. Journal of Cancer Research and Clinical Oncology, 139(5):739–746, May 2013. ISSN 1432-1335. doi: 10.1007/s00432-013-1376-6.

[50] Rex Ying, Dylan Bourgeois, Jiaxuan You, Marinka Zitnik, and Jure Leskovec. GNNExplainer: Generating Explanations for Graph Neural Networks, November 2019.

[51] Shijiao Zhi, Chen Chen, Hanlin Huang, Zhengfu Zhang, Fancai Zeng, and Shujun Zhang. Hypoxia-inducible factor in breast cancer: Role and target for breast cancer treatment. Frontiers in Immunology, 15:1370800, 2024. ISSN 1664-3224. doi: 10.3389/fimmu.2024.1370800.

[52] Yangyan Zhong, Boni Ding, Liyuan Qian, Wei Wu, and Yanguang Wen. Hormone Receptor Expression on Endocrine Therapy in Patients with Breast Cancer: A Meta-Analysis. The American Surgeon, 88(1):48–57, January 2022. ISSN 1555-9823. doi: 10.1177/0003134820972327.

