## Supplementary Table for "SpaCEy: Discovery of Functional Spatial Tissue Patterns by Association with Clinical Features Using Explainable Graph Neural Networks"

### Supplementary Material

#### *Supplementary Table 1. LUAD Dataset Statistics*

| Description | Value |
|---|---|
| Standard deviation of cells per patient | 1136.03 |
| Average cells per sample | 3996 |
| Median cells per sample | 4057 |
| Minimum cells per sample | 610 |
| Maximum cells per image | 8869 |
| Total patients in clinical data | 416 |
| Patients with survival data | 416 |
| Patients with death information | 416 |
| Patients with progression information | 404 |
| Patients with stage information | 415 |
| Patients with histological pattern information | 416 |

***Supplementary Table 2. JacksonFischer Dataset Statistics***

| **Description** | **Value** |
|---|---|
| Total number of patients | 357 |
| Total number of images | 735 |
| Average cells per image | 1730.29 |
| Standard deviation of cells per image | 1296.07 |
| Maximum cells per image | 6908 |
| Minimum cells per sample | 63 |
| Maximum cells per sample | 13291 |
| Minimum images per patient | 1 |
| Maximum images per patient | 7 |

***Supplementary table 3. METABRIC Dataset Statistics***

| Metric | Value |
|---|---|
| Total number of images | 460 |
| Total number of patients | 405 |
| Mean Images per Patient | 1.135802469 |
| Max Images per Patient | 3 |
| Std Images per Patient | 0.383866675 |
| Total number of patients | 405 |
| Total number of images | 460 |
| Average cells per image | 884.1 |
| Standard deviation of cells per image | 503.6097397 |
| Minimum cells per image | 7 |
| Maximum cells per image | 3085 |
| Minimum cells per sample | 7 |
| Maximum cells per sample | 3438 |
| Minimum images per patient | 1 |
| Maximum images per patient | 3 |
| Median Images per Patient | 1 |
| Std Images per Patient | 0.383866675 |

***Supplementary Figure S1. Distribution of clinical subtypes across clusters in the JacksonFischer dataset.***

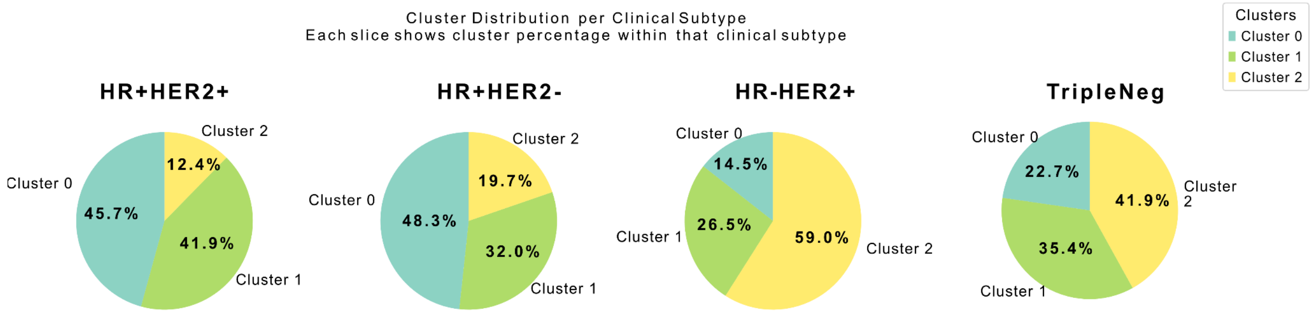

***Supplementary table 4. Baseline Hyperparameters***

| Model | Hyperparameter | Values / Range |
|---|---|---|
| **Linear Regression** | fit_intercept | [True, False] |
| | positive | [True, False] |
| **Support Vector Machine** | kernel | ["poly", "rbf", "sigmoid"] |
| | C | linspace(0.1, 50, 100) |
| | gamma | [10^x for x in -5 to 4] |
| | max_iter | range(50, 200) |
| **Random Forest** | n_estimators | linspace(10, 50, 40) |
| | max_features | ["auto", "sqrt", "log2"] |
| | bootstrap | [True, False] |
| | max_depth | range(1, 32) |
| | min_samples_split | linspace(0.1, 1.0, 9) |
| **Multilayer Perceptron (MLP)** | hidden_layer_sizes | ([100–299], [1–4]) |
| | activation | ['logistic', 'tanh', 'relu'] |
| | solver | ['sgd', 'adam'] |
| | alpha | linspace(0.0001, 0.001, 10) |
| | batch_size | [100, 150] |
| | learning_rate | ['constant', 'invscaling', 'adaptive'] |
| | max_iter | range(20, 50) |
| **Decision Tree** | criterion | ['mse', 'friedman_mse', 'mae'] |
| | max_features | ["auto", "sqrt", "log2"] |
| | max_depth | range(1, 32) |
| | min_samples_split | linspace(0.1, 1.0, 9) |
| | max_leaf_nodes | range(2, 20) |
| **Gradient Boosting** | loss | ['ls', 'lad', 'huber', 'quantile'] |
| | learning_rate | linspace(0.01, 0.15, 10) |
| | n_estimators | range(75, 150) |

| | | |
|---|---|---|
| | criterion | ['mse', 'mae', 'friedman_mse'] |
| | max_depth | range(3, 10) |
| | subsample | linspace(0.5, 1.5, 10) |
| **FastSurvivalSVM** | max_iter | [20, 100, 500, 1000] |
| | tol | [1e-1, 1e-2, 1e-3, 1e-5] |
| | optimizer | ['avltree', 'rbtree', 'simple'] |
| **RandomSurvivalForest** | n_estimators | range(100, 1001, 100) |
| | min_samples_split | [5, 10, 20, 30] |
| | min_samples_leaf | [10, 15, 30] |
| **GradientBoostingSurvivalAnalysis** | n_estimators | range(100, 1001, 100) |
| | learning_rate | linspace(0.1, 1.0, 10) |
| | max_depth | range(1, 4) |

***Supplementary Table 5. Scalability Analysis***

| Num of markers | Number of Samples | Average Nodes Per Sample | Average Epoch Time | Total Training Tİme | Peak GPU Memory | Average GPU Memory Utilizatior |
|---|---|---|---|---|---|---|
| 30 | 50 | 1221.94 | 1.61605193 | 16.1605194 | 26.91210 | 99.8 |
| 30 | 100 | 1252.86 | 1.23026197 | 12.3026197 | 32.31298 | 90.5 |
| 30 | 200 | 1246.83 | 4.23998074 | 42.3998075 | 40.78027 | 100 |
| 30 | 500 | 1282.726 | 8.33640127 | 83.3640127 | 72.40332 | 91.5 |
| 30 | 1000 | 1297.171 | 9.12987425 | 91.2987425 | 68.21582 | 88.7 |
| 30 | 1500 | 1290.31 | 15.3915977 | 153.915977 | 66.07910 | 91.2 |
| 50 | 50 | 1221.94 | 0.69628000 | 6.96280003 | 28.84277 | 91.8 |
| 50 | 100 | 1252.86 | 1.38614032 | 13.8614032 | 29.92626 | 91.7 |
| 50 | 200 | 1246.83 | 2.46376423 | 24.6376424 | 41.06396 | 91.8 |
| 50 | 500 | 1282.726 | 5.77550354 | 57.7550354 | 71.87841 | 91.2 |
| 50 | 1000 | 1297.171 | 10.9450917 | 109.450918 | 79.34326 | 91.2 |
| 50 | 1500 | 1290.31 | 12.7633330 | 127.633331 | 68.49121 | 89.6 |
| 100 | 50 | 1221.94 | 0.62680566 | 6.26805663 | 28.69677 | 89.9 |
| 100 | 100 | 1252.86 | 1.36121165 | 13.6121166 | 32.39257813 | 89.5 |
| 100 | 200 | 1246.83 | 3.10268073 | 31.0268073 | 47.12353516 | 99.3 |
| 100 | 500 | 1282.726 | 9.11942162 | 91.1942163 | 78.5078125 | 91.3 |
| 100 | 1000 | 1297.171 | 17.7751340 | 177.751341 | 79.22021484 | 90 |
| 100 | 1500 | 1290.31 | 22.8611951 | 228.611952 | 73.33691406 | 89.9 |
| 500 | 50 | 1221.94 | 1.16837935 | 11.6837935 | 39.04931641 | 99.8 |
| 500 | 100 | 1252.86 | 2.14238290 | 21.4238291 | 47.328125 | 99.7 |
| 500 | 200 | 1246.83 | 3.74223072 | 37.4223073 | 77.26855469 | 99.8 |
| 500 | 500 | 1282.726 | 8.84009161 | 88.4009161 | 149.7998047 | 89.8 |
| 500 | 1000 | 1297.171 | 9.23399851 | 92.3399851 | 138.465332 | 85.6 |
| 500 | 1500 | 1290.31 | 15.8711196 | 158.711197 | 133.4165039 | 90.1 |
| 1000 | 50 | 1221.94 | 0.64391703 | 6.43917036 | 51.85107422 | 92 |
| 1000 | 100 | 1252.86 | 1.475946 | 14.7594619 | 78.89257813 | 91.8 |

| | | | | | | |
|---|---|---|---|---|---|---|
| 1000 | 200 | 1246.83 | 2.42435278 | 24.2435279 | 112.2890625 | 91.6 |
| 1000 | 500 | 1282.726 | 5.89035170 | 58.903517 | 235.1777344 | 90.9 |
| 1000 | 1000 | 1297.171 | 10.9387927 | 109.387927 | 222.7954102 | 88.1 |
| 1000 | 1500 | 1290.31 | 15.5928630 | 155.92863 | 200.1323242 | 87.4 |

***Supplementary Table 6. Hyperparameter values for model training***

| Category | Hyperparameter | Values | Description |
|---|---|---|---|
| **Model Architecture** | Number of GCN Layers | 2 | Number of graph convolutional layers |
| **Model Architecture** | Number of Feed-Forward Layers | 1, 2 | Number of fully connected layers |
| **Model Architecture** | GCN Hidden Dimension | 16, 32, 64, 128 | Hidden dimension size for graph convolutional layers |
| **Model Architecture** | Fully Connected Layer Dimension | 64, 128, 256 | Hidden dimension size for fully connected layers |
| **Model Architecture** | Dropout Rate | 0.2, 0.3 | Dropout probability for regularization |
| **Training Parameters** | Learning Rate | 0.1, 0.01, 0.001, 0.0001 | Initial learning rate for optimization |
| **Training Parameters** | Batch Size | 16, 32, 64 | Number of samples per training batch |
| **Training Parameters** | Epochs | 200 | Maximum number of training epochs |
| **Training Parameters** | Weight Decay | 0.1, 0.001, 0.0001, 1e-5 | L2 regularization coefficient |
| **Learning Rate Scheduler** | Factor | 0.5, 0.8, 0.2 | Factor by which learning rate is reduced |
| **Learning Rate Scheduler** | Patience | 5, 10 | Number of epochs with no improvement before reducing LR |
| **Learning Rate Scheduler** | Minimum Learning Rate | 0.00002, 0.0001 | Lower bound for learning rate |
| **Model-Specific** | GAT Heads | 1, 3, 5 | Number of attention heads (GAT model only) |
| **Model-Specific** | PNA Aggregators | min, max, sum, mean, sum max | Aggregation functions (PNA model only) |
| **Model-Specific** | PNA Scalers | identity, amplification | Scaling functions (PNA model only) |

*This table lists the key hyperparameters and their respective search ranges explored during model optimization. Each model was tuned using grid or randomized search strategies within these parameter spaces to identify optimal configurations for predictive performance and generalization. The hyperparameter ranges include structural, regularization, and training parameters relevant to both classical machine learning and survival analysis models.*

***Supplementary Figure S2. GNNExplainer model training***

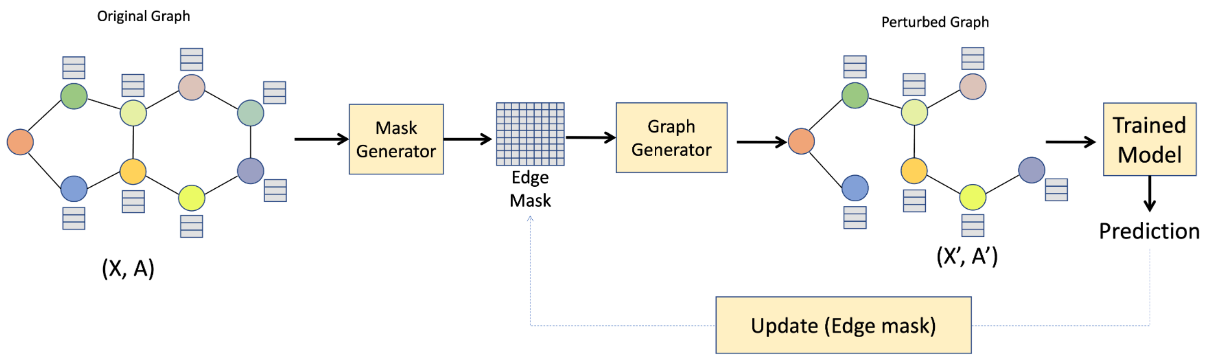


The general framework of GNNExplainer considers a graph G=(X,A), where the input graph is first passed through a Mask Generator. The masked representations are then processed by a Graph Generator to project them back into the original graph space, yielding(X′,A′). These modified inputs are fed into the trained GNN model and the resulting prediction f(X′,A′) is used in a mutual information–based loss function to optimize the Mask Generator. The optimization process ultimately produces refined masks that highlight the most relevant nodes, and edges for the model's prediction.
